# Contrastive Regulatory Embeddings Attention Model for Differential Expression Prediction with Personalized Genomes: Insights and Challenges

**DOI:** 10.64898/2026.08.02.742309

**Authors:** Zhirui Hu, Jason Ku, Katherine Pollard

## Abstract

Deep learning models applied to DNA sequences have achieved success in predicting gene expression, chromatin profiles, and variant pathogenicity. However, learning cross-individual differences remains challenging because DNA sequence variation between individuals is small, while gene expression is heavily influenced by non-genetic noise. Prior efforts to predict personalized gene expression from sequence have shown limited generalizability beyond training genes, revealing limitations such as the dilution of variant signals among highly similar input sequences across consecutive convolutional downsampling layers. In this work, we explore architectural modifications addressing these challenges. We propose a contrastive regulatory embedding attention model (CREAM) designed to better capture subtle sequence differences between individuals. To mitigate the impact of non-genetic variability, we decompose gene expression into genetic and non-genetic components and evaluate model performance on the genetic signal. Evaluated on simulated and GTEx transcriptomic datasets, CREAM captures tissue-specific gene expression and significantly outperforms baseline architectures on training genes. CREAM autonomously prioritizes statistically fine-mapped causal expression quantitative trait loci, and introducing an auxiliary ***L*_1_** inductive bias further sharpens localizing causal variant in unseen genes. However, generalization to predicting expression of unseen genes collapses to near-zero correlation among all tested methods. Single-run model predictive uncertainty capture prediction accuracy in training genes and mirrors cross-run consistency in test genes. While accurate inference on unseen genes remains an open problem, our results highlight key obstacles and suggest directions for modeling personalized gene regulation from DNA sequence.

## 1 Introduction

Deciphering the functional code embedded within non-coding DNA represents a fundamental challenge in human genetics and molecular biology. The standard approach to linking non-coding genetic variants to transcription has relied on regularized linear modeling frameworks, such as PrediXcan [1]. These associated variants are called expression quantitative trait loci (eQTL). Although these linear models are robust and explain a substantial portion of expression heritability, they suffer from fundamental limitations. Because they are parameterized on predefined variants per gene, linear models cannot generalize to rare or de novo variants, nor can they model variants in multiple genes together to identify shared regulatory mechanisms. Furthermore, these frameworks fail to provide a mechanistic interpretation of highly correlated associated variants because they do not account for the local sequence context, such as the presence or disruption of transcription factor (TF) binding motifs.

Deep learning architectures applied to DNA sequences have achieved remarkable success in decoding the complex regulatory grammar of the genome. By mapping raw sequence inputs to genome-wide epigenomic features, sequence-to-function (S2F) models have learned regulatory grammars and enabled highly accurate predictions of cell type-specific gene expression, chromatin accessibility profiles, and variant pathogenicity. By leveraging deep convolutional architectures, models such as Basset[2] and Basenji[3] successfully capture localized features, such as TF binding motifs. The integration of transformer blocks and U-Net, e.g. Enformer[4] and Borzoi[5] extended the receptive field to nearly 200kb and 500kb, allowing the model to capture longer sequence context and predicting tissue-specific gene expression measured by RNA sequencing. Together, these reference-trained models have achieved unprecedented accuracy in predicting cross-gene expression levels and functional genomic profiles across held-out chromosomes. However, these foundational models are predominantly trained and evaluated on a single, invariant reference genome, but not sequence variations across individuals. Consequently, adapting these architectures to decode personal DNA sequences and capture cross-individual expression dynamics remains a major bottleneck.

Zero-shot predicting personalized gene expression directly from individual-level genomes encounters both biological and computational challenges. First, because reference-trained deep learning models have never observed genetic polymorphisms during their training phase, they face limitations in predicting cross-individual expression differences [6]. Second, the genomic sequence variation between any two individuals is minor, comprising sparse single-nucleotide variants (SNVs) and small indels within mostly identical sequence background, whereas individual molecular profiles, e.g. gene expression, are confounded by non-genetic and environmental noise [7].

Recent benchmarking studies indicate that deep learning-based DNA sequence models consistently struggle to outperform traditional linear models, such as regularized elastic net models in mapping eQTLs, when predicting expression levels across a population [6, 7]. A benchmark study on four state-of-the-art architectures, i.e. Enformer, Basenji2 [8], ExPecto [9], and Xpresso [10], on paired whole-genome sequencing (WGS) and RNA-seq data from the Geuvadis consortium [11], revealed that while these models explain a large fraction of the variance in expression across different genes in a single reference individual, their predictive power degrades to near zero when predicting the expression of a single gene across different individuals. Aside from successfully identifying the large phenotypic effects of rare and disruptive variants [12], current deep sequence models demonstrate poor generalizability when predicting individual-specific expression for genes outside the training set [6].

Systematic evaluations have brought to light critical structural pitfalls within current deep sequence models that explain the lack of predictive sensitivity. Notably, state-of-the-art models regularly fail to correctly predict the direction of variant effects, a flaw primarily attributed to an insufficiently learned sequence motif grammar [13]. Furthermore, standard S2F models employ consecutive deep convolutional layers that progressively downsample sequences to compress long-range context (e.g., aggregating features into 128 bp bins in Enformer). This reduction in spatial resolution frequently dilutes fine-scale nucleotide differences between individuals, making it difficult for downstream layers to discern the impact of individual variants.

Beyond sequence resolution loss and poor generalized performance, standard S2F deep learning models do not provide predictive confidence. Quantifying predictive confidence is essential for accurately interpreting variant effects, guiding downstream experimental validation, and advancing the development of personalized medicine.

Establishing reliable uncertainty quantification (UQ) has shown exceptionally difficult. Recent benchmark evaluations have revealed calibration anomalies in state-of-the-art deep sequence models: networks routinely yield high-confidence predictions on standard reference genomes even when the predicted expression is incorrect, yet conversely produce highly inconsistent or low-confidence predictions when evaluated on sequences containing expression quantitative trait loci (eQTLs). Across model replicates optimized with identical data but different random initializations, variant effect predictions have been shown to diverge or switch signs in more than half of evaluated cases [14]. While recent frameworks have attempted to resolve these discrepancies by applying deep ensembles and alternative regression metrics to baseline models [15], or utilizing knowledge distillation techniques [16] to estimate epistemic variation, current models applied to cross-individual predictions remain inadequate to explicitly detect when a personalized variant prediction is not reliable. Consequently, integrating robust, self-aware uncertainty intervals is a necessity to prevent high confidence but false positive variant interpretations in personalized medicine.

To address high sequence similarity, spatial resolution loss, and poor generalization to unseen genes, we explore structural modifications designed to resolve these pitfalls. We propose the Contrastive Regulatory Embedding Attention Model (CREAM), an architecture to better isolate cross-individual genetic variation and amplify causal eQTL signals. CREAM introduces a contrastive mechanism that identifies non-identical multi-scale genomic bins between a pair of individual sequences, and contextualizes these localized variants against global sequence backgrounds through attention layers. To retain multi-scale resolution and mitigate spatial coarse-graining, our framework establishes direct links from intermediate convolutional layers directly to the attention layers. Additionally, we incorporate prior biological knowledge by utilizing known expression quantitative trait loci (eQTLs) to guide the attention weights, explicitly biasing the model toward curated regulatory variants.

Recognizing that non-genetic variation in gene expression can mask genetic signals and degrade predictive performance, we explicitly decompose individual gene expression into genetic and non-genetic components, evaluating our model directly on the genetic signal. Finally, we explore the model’s capacity to quantify predictive uncertainty, assessing if the model can accurately detect its own out-of-sample prediction errors. Using Whole Genome Sequencing (WGS) and paired transcriptomic data from three tissues in the Genotype-Tissue Expression (GTEx) V8 dataset, we fine-tune Enformer backbone and additional model layers in CREAM. Our results demonstrate that while gene generalization challenges remain, CREAM successfully uncovers tissue-specific eQTLs and introduces a principled framework for personalized gene regulation modeling.

## 2 Results

### 2.1 Overview of CREAM architecture and datasets

To capture the subtle regulatory impact of individual genomic variations, we developed the Contrastive Regulatory Embedding Attention Model (CREAM), using the S2F model Enformer as the backbone. Enformer is suboptimal for to resolve minor cross-individual sequence variations because nearby variants become indistinguishable in the final embedding layer (Figure A1b). Furthermore, because the effects of individual SNVs are frequently flattened by successive downsampling layers, simply fine-tuning Enformer to use the embedding of a focused bin is inadequate to capture the variant’s impact (Figure A1a). To overcome this limitation, we modified the model structure to explicitly isolate sequence variations, incorporate genomic contexts and reframed the training objective to predict relative differential expression between pairs of individuals rather than absolute levels, which removes the dominant gene-level baseline.

The CREAM framework operates through a multiscale feature retrieval and cross-attention mechanism [17], detailed as follows (Figure 1):

1) Multiscale genomic features: to circumvent the loss of spatial resolution in Enformer’s deeper convolution layers, CREAM establishes direct connections from intermediate layers to its final custom attention mechanism. Specifically, the framework identifies sequence bins at each convolutional layer that are polymorphic (non-identical) between the paired input sequences of two individuals. It then extracts and concatenates the multi-resolution embeddings from these variant-containing bins alongside those containing the transcription start site (TSS). This multi-scale concatenation preserves high-resolution sequence contexts for SNVs.
2) Contrastive cross-attention: rather than processing highly similar sequences independently, CREAM implements a contrastive cross-attention module. Using the multi-scale embeddings of the variant and TSS from the first step, the query vector (Q) is constructed by further concatenating the embeddings from both input sequences. Concurrently, keys (K) and values (V) are generated by concatenating the representations from Enformer’s final attention layer for both sequences, capturing the surrounding genomic background. This formulation allows the attention mechanism to dynamically transform localized variant queries by contextualizing them within the global sequence background.
3) Attention pooling and variant interpretation: to aggregate the polymorphic signals scattered across distinct loci, we employ an attention-pooling layer that dynamically weights and summarizes the joint effects of all cross-individual SNVs and the TSS into a unified, low-dimensional embedding. This pooled representation is subsequently passed to a Multi-Layer Perceptron (MLP) to predict gene expression differences. Crucially, the learned attention weights provide direct model interpretability by reflecting the relative regulatory contribution of local genomic bin around each individual SNV, allowing for direct validation against posterior inclusion probabilities (PIPs) from statistical fine-mapping tools like SuSiE [18, 19]. This architecture parallels recent shifts toward attention-based sequence frameworks capable of capturing long-range regulatory interactions for variant interpretation [4].
4) Prior-knowledge eQTL guidance: to insulate the model from extensive transcriptomic noise and prevent it from overfitting to background non-genetic signals, we implement an optional inductive bias during training. We introduce an auxiliary loss function that minimizes the L1 distance between the model’s learned attention weights and the effect sizes of known eQTLs. This optimization constraint forces the network to prioritize curated regulatory variants over non-causal background variations.

**Fig. 1:**
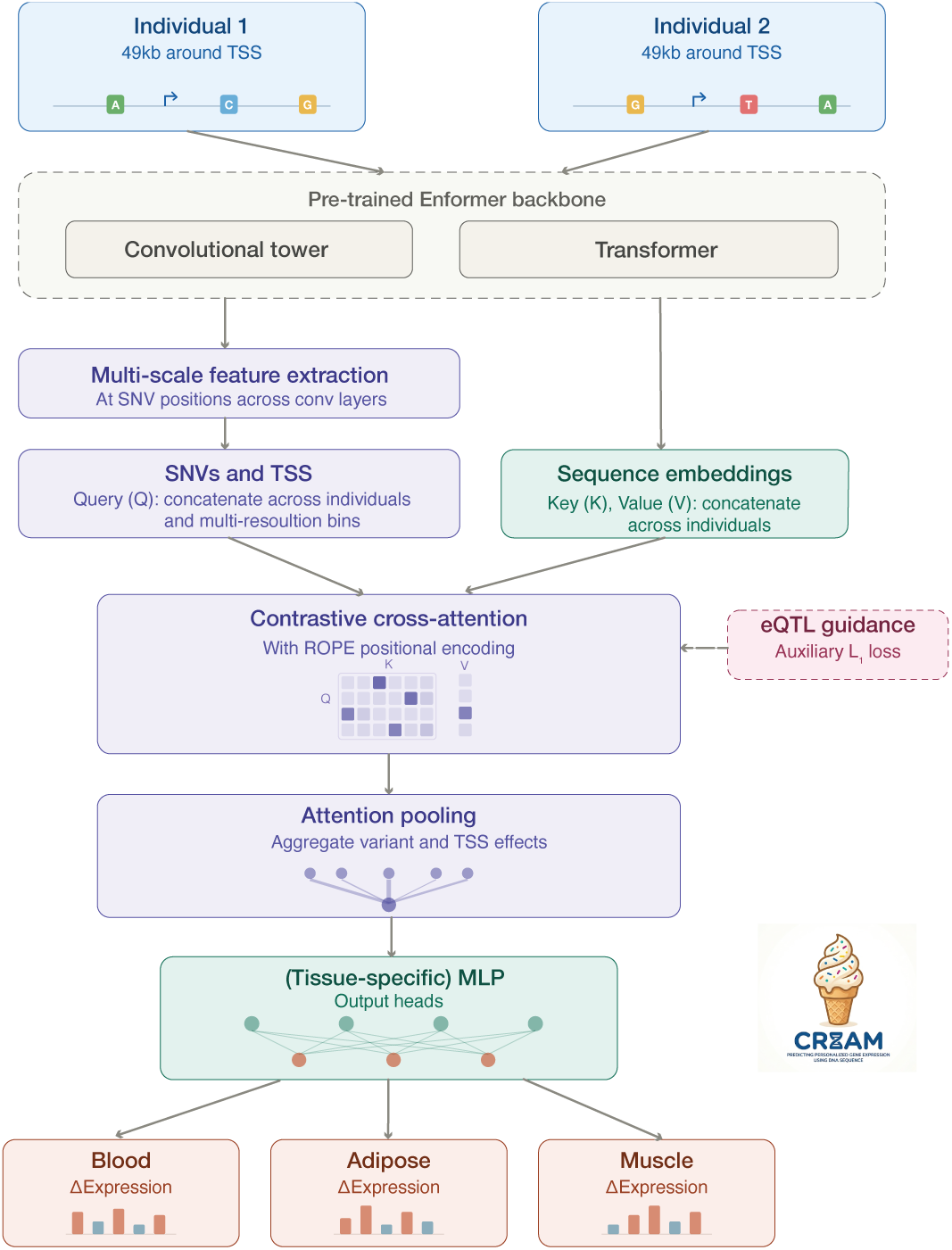
Schematic illustrations of CREAM architecture. Input and Pre-trained Backbone: Paired genomic sequences (49 kb centered on the TSS) from two distinct individuals are processed through a pre-trained Enformer backbone. Multi-scale Feature Extraction for Query (*Q*): Intermediate convolutional layers are scanned to extract multiscale features at polymorphic (SNV) bins and the TSS, which are concatenated across layers and both individuals to form *Q*. Keys (*K*) and Values (*V*): Full sequence embeddings across all genomic bins are extracted from Enformer’s transformer attention layers and concatenated across both individuals to form *K* and *V*. Contrastive Cross-Attention and eQTL Guidance: Localized variant queries (*Q*) are mapped against the global regulatory environment (*K, V*) using cross-attention equipped with Rotary Position Embedding (RoPE). Optionally, attention weights are constrained during training using an auxiliary *L*_1_ loss guided by known eQTL effect sizes. Attention Pooling and Multi-Tissue Output: An attention-pooling layer aggregates spatially dispersed variant and TSS effects into a unified representation. Downstream MLP output heads (configurable as shared or tissue-specific) project this representation to predict cross-individual differential expression (ΔExpression) across target tissues (e.g., blood, adipose, muscle).

Because eQTL regulatory mechanisms can be either highly tissue-specific or widely shared across human tissues, we explored three distinct structural variations of CREAM to optimize multi-tissue expression predictions: CREAM 1 (Tissue-specific attention and shared output), which utilizes separate attention layers for each tissue to capture distinct, tissue-dependent eQTLs, while passing the pooled embeddings into a single, shared output layer; CREAM 2 (Fully decoupled attention and output), that employs both separate attention layers and independent tissue-specific output layers, allowing complete decoupled parameters across distinct tissues to capture independent regulatory programs; CREAM 3 (Shared attention and tissue-specific output), which constrains the model to use a single, shared attention layer across all tissues, relying entirely on tissue-specific downstream output layers to resolve expression differences. We evaluated our framework using the Genotype-Tissue Expression (GTEx) V8 dataset [20], leveraging its paired whole-genome sequencing (WGS) and transcriptomic profiles across individuals. To rigorously assess model generalizability, we implemented a dual-partitioning scheme, splitting both the individuals (donors) and the genes into distinct training, validation, and test sets. We filtered for genes that exhibited a minimum expression variance and possessed at least one causal eQTL in at least one of the three target tissues (see Methods).

Within a 49 kb window of the TSS, we obtained around 50K causal eQTLs per tissue genome-wide, and around 120K unique eQTLs in total. Most genes harbor only a handful of causal eQTLs, with a median of 8, highlighting the inherent difficulty of identifying these sparse regulatory variants. Most identified eQTLs possess low PIP values (*<* 0.2) while only a small fraction approach 1.0, confirming that widespread collinearity among proximal SNVs restricts the capacity of standard linear models to pinpoint causal loci (Figure A2).

### 2.2 Prediction performance

To validate the capability of CREAM to capture true genetic effects independently of environmental or technical confounding, we first evaluated the model using a controlled simulation framework then on real GTEx data. We leveraged fine-mapping outputs from SuSiE applied to the GTEx cohort across three primary tissues: whole blood, adipose, and skeletal muscle. These tissues were selected due to their large sample sizes and high abundance of eQTLs. Treating these statistically fine-mapped eQTLs as biological ground truths, we utilized their PIPs and effect sizes as coefficients within an additive linear model. By combining these coefficients with individual genotypes, we simulated personalized, noise-free gene expression profiles to provide a rigorous benchmark for cross-individual prediction.

We benchmarked CREAM models against alternative frameworks including the baseline model, which directly fine-tunes Enformer on the training data, and the contrast model, which leverages a contrastive embedding and cross-attention between TSS bin and all sequence bins, regardless of sequence variations (see Methods).

#### 2.2.1 Simulations

As expected, in the noise-free simulations, when evaluating training genes across unseen test donors, our model exhibits near-perfect performance, with both Pearson’s *r* and coefficient of determination (*R*^2^) approaching 1.0 (Figure 2a, 2b). Because causal eQTLs identified by SuSiE constitute common variants, test donors do not have novel polymorphisms whose regulatory effects were completely omitted during training. Furthermore, the capacity of the model is sufficiently large to parameterize all eQTL effects across the training gene set.

**Fig. 2:**
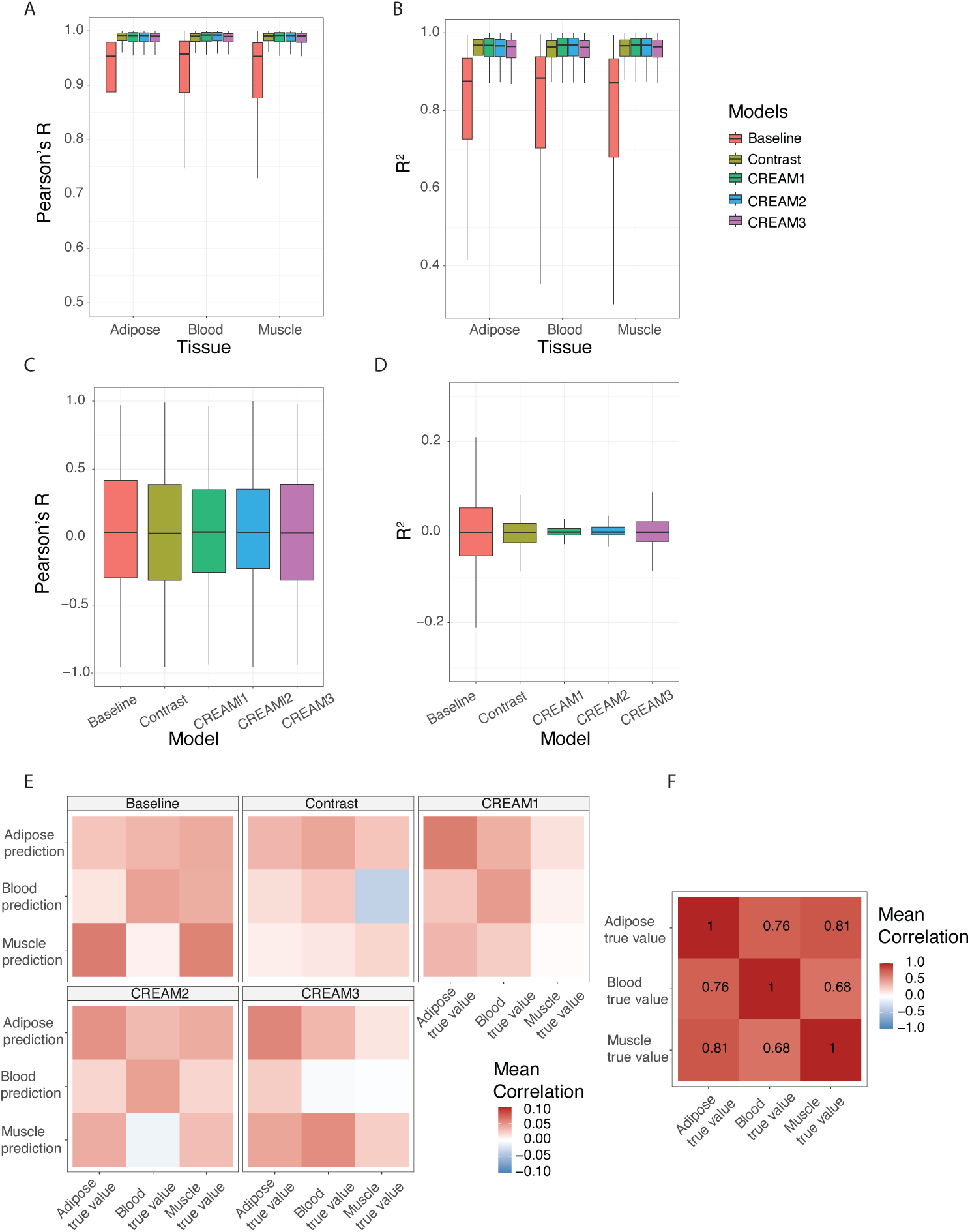
Model performance in simulation study. (A) and (B) CREAM outperforms baselines in training genes. The boxplot shows the distribution of (A) Pearson’s R and (B) R2 (Y-axis) among training genes evaluated across test donors. X-axis shows different tissues. (C) and (D) CREAM has similar performance as baselines in test genes. The boxplot shows the distribution of (C) Pearson’s R and (D) R2 (Y-axis) among test genes evaluated across test donors. X-axis shows different models. For (A) - (D), colors represent different models. CREAM 1-3 are variant_1_s_5_of CREAM. (E) Prediction of CREAM 2 has better tissue-specificity for test genes. Cross-tissue correlation of predicted and true gene expression in different models. Each row is predicted gene expression in different tissues and each column is true simulated gene expression. Each panel shows the result from an evaluated model. (F) Cross-tissue correlation of true simulated gene expression. Both rows and columns are true simulated gene expression.

We observed minor performance variations among the different architectural configurations. CREAM 1 and CREAM 2, which have tissue-specific attention blocks, slightly outperform the alternative structure (CREAM 3), suggesting that decoupling the attention layers across tissues is critical to accommodate tissue-specific eQTL profiles and distinct regulatory mechanisms (Figure 4e). On the other hand, once these variable eQTL effects are aggregated into a unified embedding, a shared downstream network is sufficient to predict final gene expression differences in multiple tissues (i.e., CREAM 2’s performance is similar to CREAM 1). Compared with alternative frameworks, CREAM consistently outperforms the baseline model and marginalizes the contrast model. These performance trends remain highly robust across independent training seeds and donor cross-validation splits (Figure A3, Figure A4).

However, for unseen test genes, CREAM’s predictive performance remains limited despite the extensive structural modifications to the neural network architecture (Figure 2c, 2d). The median performance metrics are around zero across all evaluated models, which suggests that the regulatory mechanism learned from training genes do not readily generalize to out-of-sample test genes. Nevertheless, when evaluating the cross-tissue correlation between predicted and true simulated gene expression, CREAM 2 demonstrates superior capacity to capture tissue-specific expression dynamics compared to alternative architectures (Figure 2e). Specifically, CREAM 2 yields both higher diagonal correlation coefficients between matching predicted and true tissues, and captures expected off-diagonal biological signatures, such as the elevated cross-correlation between the true and the predicted expression in adipose and muscle tissue respectively, reflecting their true gene expression profiles (Figure 2f). These trends remain robust when evaluating subsets of tissue-specific genes (Figure A5). Taken together, these findings indicate that implementing fully decoupled, tissue-specific attention blocks and downstream output layers is structurally essential for accurately resolving divergent gene expression patterns across distinct tissues.

#### 2.2.2 GTEx data

We evaluated the predictive performance of the baseline, contrast, and CREAM 2 architectures using standardized gene expression data from the GTEx dataset across blood, adipose, and muscle tissues. To ensure consistency and direct comparability, we maintained the identical training/test donor splits and gene partitions established in our simulation framework. For the training genes, both the Contrast model and CREAM consistently outperform the baseline architecture across all three tissues (Figure 3a, 3b). The median Pearson correlation coefficients (*r*) center around 0.1, which is probably due to the substantial non-genetic and environmental noise inherent to empirical transcriptomic data. Cross-individual per gene prediction performance is essentially unchanged for held-out donors carrying de novo SNVs (Figure A6), indicating that such variants are not a major driver of the limited predictive accuracy.

**Fig. 3:**
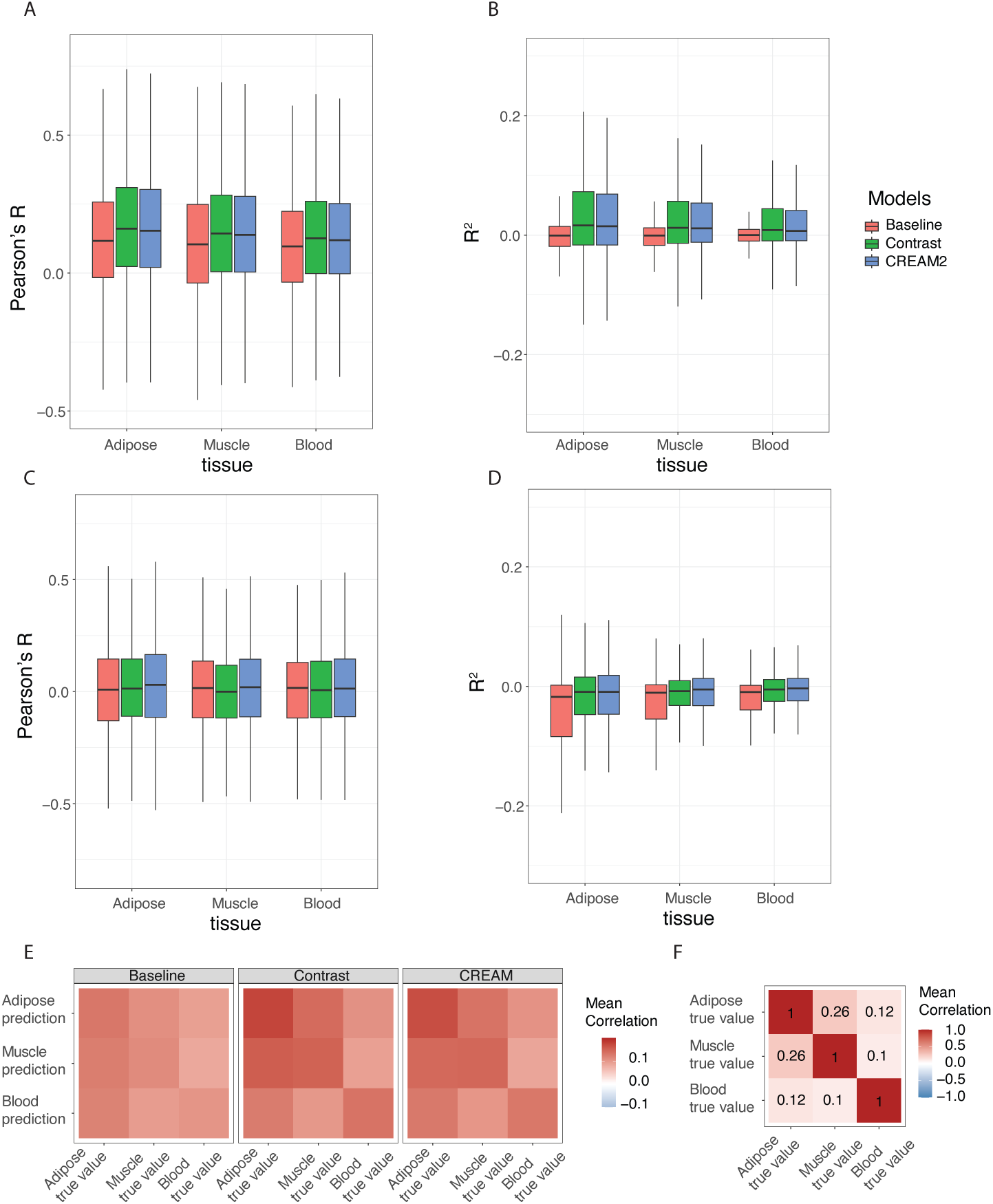
Model performance in GTEx data. (A) and (B) CREAM 2 outperforms baselines in training genes. The boxplot shows the distribution of (A) Pearson’s R and (B) R2 (Y-axis) among training genes evaluated across test donors. (C) and (D) CREAM 2 has similar performance as baselines in test genes. The boxplot shows the distribution of (C) Pearson’s R and (D) R2 (Y-axis) among test genes evaluated across test donors. For (A) - (D), colors represent different models. X-axis represents different tissues. (E) Prediction of CREAM 2 has better tissue-specificity for test genes. Cross-tissue correlation of predicted and true gene expression from different models. Each row is predicted gene expression in different tissues and each column is true gene expression. (F) Cross-tissue correlation of true gene expression. Both rows and columns are true gene expression.

Despite these modest correlation magnitudes, CREAM 2 successfully captures tissue-specific gene expression patterns (Figure 3e). Specifically, tissue-specific predictions generated by CREAM 2 exhibit higher correlation with the true observed values in their corresponding target tissues. In contrast, predictions from the baseline model suffer from systemic bias, remaining disproportionately correlated with adipose gene expression across all evaluated tissues. These tissue-specific trends remain robust when the analysis is restricted exclusively to tissue-specific genes (Figure A7).

However, the poor unseen gene generalization remains a major bottleneck for all evaluated methods. This indicates that the regulatory features learned from the training genes fail to generalize to unseen test genes, resulting in unseen gene prediction performance that falls near zero (Figure 3c, 3d).

### 2.3 Attention weights capture tissue-specific eQTLs

#### 2.3.1 Simulations

Next, we evaluated the model’s interpretability by comparing the learned attention weights between causal eQTLs and other SNVs. For the training genes, the architecture consistently allocates higher attention weights to fine-mapped eQTLs (PIP *>* 0.7) relative to background SNVs (Figure 4a), demonstrating that the model is capable of autonomously identifying causal variants driving gene expression. We also observed that CREAM preferentially weights causal SNVs in unseen test genes, albeit to a lesser extent (Figure 4b). We hypothesize that gene regulation is a complex, multi-scale process involving parallel and combinatorial pathways. Consequently, while the model successfully generalizes its attention to casual eQTLs of unseen genes, prioritizing these SNVs remains insufficient for precise downstream gene expression prediction.

**Fig. 4:**
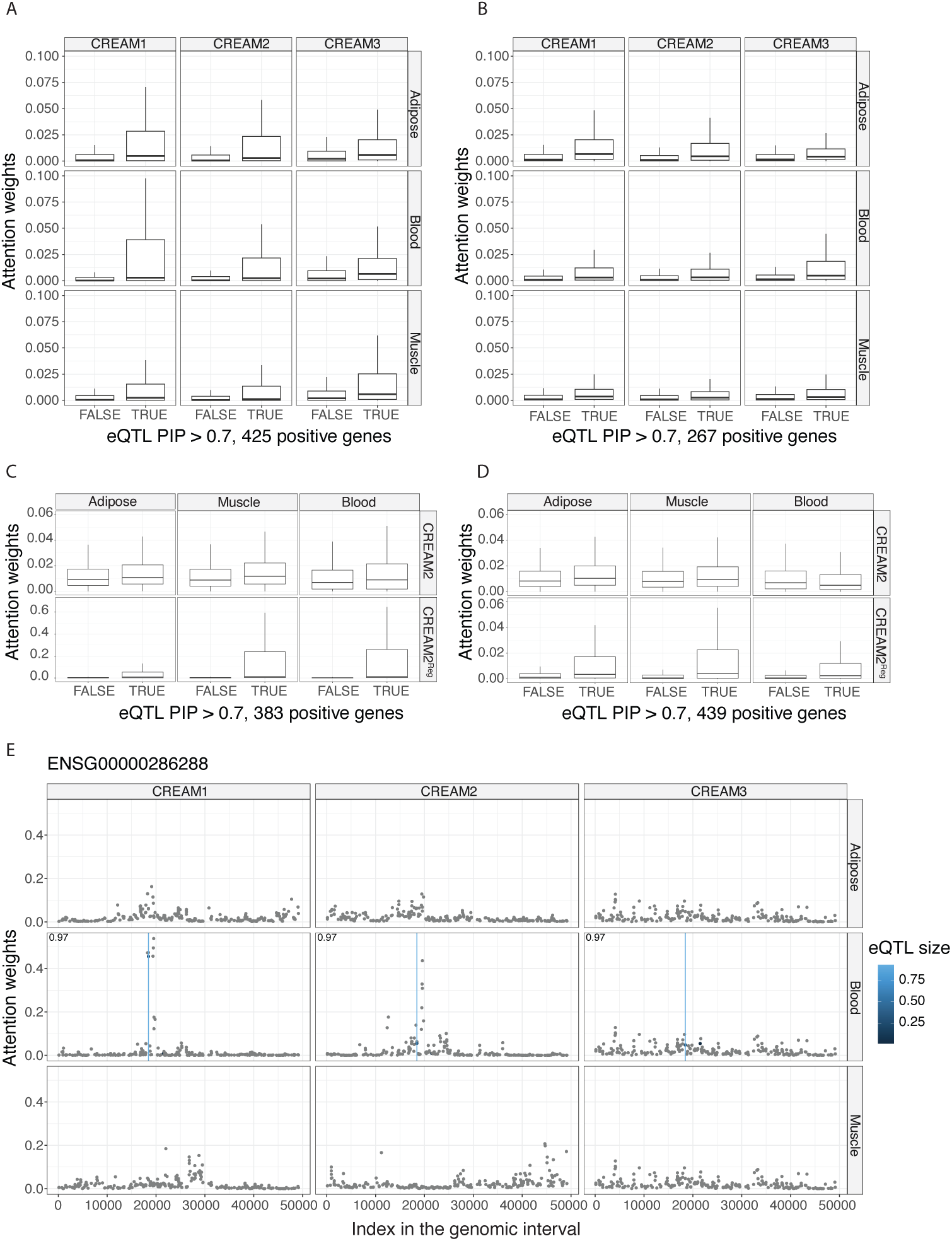
Attention weights in simulation and GTEx data. (A)-(B) show the distribution of attention weights of each SNV (Y-axis) on 425 selected training genes (A) and selected 267 test genes (B) in simulated data. Each column is a CREAM model and each row is a tissue. (C)-(D) show the distribution of attention weights of each SNV (Y-axis) on 383 selected training genes (C) and selected 439 test genes (D) in GTEx data. Each column is a tissue. Top row is a CREAM 2 and bottom row is CREAM 2 with eQTL guided attention weights (CREAM2*^Reg^*). (A) - (D) We select genes with at least one eQTL with PIP > 0.7, and we treat eQTL with PIP *>* 0.7 as positive samples and the rest as negative samples. Attention weights are obtained from random pairs of individuals in the test donors. As we have much more negative than positive eQTLs, we subsampled 5 times more negative than positive SNVs for each tissue. (E) Manhattan plot shows the average attention weights of each SNV in an example training gene in simulation. The attention weight is computed averaging over test donor pairs and two runs with different seeds. Y-axis is the average attention weights and X-axis is the index of input genomic interval. Each dot is a SNV colored by eQTL effect size. Vertical line indicates the eQTL with PIP *>* 0.7. Pearson’s *R* of gene expression prediction is shown in the top-left corner. Calculation is restricted to blood, as gene expression lacks variance in the remaining tissues due to a blood-specific eQTL.

Furthermore, CREAM effectively captures distinct, tissue-specific eQTL profiles (Figure 4e). For instance, gene ENSG00000286288 harbors a causal eQTL exclusive to whole blood. CREAM 1 and CREAM 2 successfully capture this variant by assigning large attention values uniquely in blood, whereas CREAM 3 fails to capture the signal, likely due to having shared attention weights across tissues. Similarly, gene ENSG00000204099 possesses distinct eQTLs in adipose and muscle tissues (Figure A8). CREAM 1 and CREAM 2 accurately align their attention profiles to these tissue-specific eQTLs within the respective tissues, whereas CREAM 3 exhibits uniform, uninformative attention weights across the locus. We observed a more dynamic regulatory landscape for gene ENSG00000249159, which contains an causal eQTL shared between adipose and muscle upstream of the TSS (Figure A8). CREAM 1 and CREAM 2 successfully concentrate higher attention upstream of the TSS in adipose and muscle tissues, yet shift their attention to downstream of TSS in blood; conversely, CREAM 3 statically places large attention weights upstream across all three tissues. Although all three models yield comparable overall predictive performance for these particular genes, CREAM 1 and CREAM 2 offer substantially greater biological interpretability via their tissue-specific attention mechanisms. We reason that CREAM 3 compensates for its shared attention layer by relying entirely on its tissue-specific downstream output layers to resolve divergent expression across tissues.

#### 2.3.2 GTEx data

Despite the substantial expression noise inherent to the empirical data, CREAM 2 consistently assigns higher attention weights to causal eQTLs than to background SNVs for training genes. This enrichment indicates that the architecture successfully learns genetic variations that contribute to personalized gene expression (Figure 4c). However, for unseen test genes, the model fails to identify causal eQTLs in certain tissues (i.e. whole blood), in contrast to the clean patterns observed in the noise-free simulation data (Figure 4d). To alleviate this generalization deficit, we introduced an auxiliary loss function during training to minimize the *L*_1_ distance between the model’s attention weights and known eQTL effect sizes. By leveraging these known effect sizes to guide the attention layers, the model, denoted as CREAM2*^Reg^*, successfully identifies causal eQTLs within unseen test genes, even when those specific variants were absent from the training set (Figure 4d). Broadly speaking, constraining the attention weights reduces background noise by driving the weights on non-eQTLs to near-zero values. While enforcing known eQTL guidance slightly penalizes performance on training genes, presumably due to its restriction of the model from learning real genetic effects missing from the curated eQTL set, it yields a slight improvement in out-of-sample generalization performance on test genes (Figure A9). Importantly, although constraining the attention weights improves the model’s detection of functional variants, this regularization does not automatically transfer to superior gene expression prediction accuracy (Figure A10). For both training and test gene sets, the eQTL-guided model more accurately detects causal eQTLs, yet its predictive performance fluctuates among genes compared to the original CREAM model (Figure A10). This variation in performance may point to complex genetic architectures not accounted for by known eQTL annotations. For example, in the case of gene ENSG00000182378 in muscle tissue, the original unconstrained CREAM model allocates higher attention weights to distal loci far from the TSS that lack known eQTL annotations. Despite this discrepancy, it achieves similar predictive accuracy to the eQTL-guided model, which prioritizes known eQTLs located adjacent to the TSS.

Furthermore, by aggregating embeddings across all sequence bins using our attention-pooling mechanism, the high-scoring variants are located further from the TSS than those highlighted by the baseline model, though they remain closer to the TSS relative to the eQTLs identified via standard linear modeling (Figure A11).

### 2.4 Uncertainty quantification

#### 2.4.1 Framework for uncertainty estimation

We evaluated the capability of the CREAM model to quantify predictive uncertainty. To achieve this, we discretized the normalized gene expression values into 10 distinct bins and modified the model’s output layer to simultaneously predict the probability distribution across these bins alongside a continuous offset from each bin’s mean value. Predictive uncertainty was then evaluated using Shannon entropy and variance metrics.

#### 2.4.2 Variations in entropy across prediction bins

The predicted entropy varied significantly across the prediction bins (Figure 5a). The central bin, representing donor pairs with near-zero or exact zero expression differences, exhibited the lowest overall entropy. This pattern is attributable to the predominance of zero-difference values in the simulated dataset (comprising 69% of the entries), which can occur when two individuals share identical genotypes at causal eQTL loci. Furthermore, certain genes naturally lack causal eQTLs in specific tissues, rendering their expression invariant across individuals. These factors produce a large pool of true zero expression differences, which the model predicts with high confidence. Conversely, the extreme peripheral bins (the leftmost and rightmost intervals) also displayed reduced entropy, indicating that the model obtains high certainty when predicting large and extreme expression differences. The model successfully learns this entropy distribution trend from the training set and generalizes it to out-of-sample test genes. The absolute predicted entropy for test genes is systematically larger, properly reflecting increased prediction uncertainty. This elevated uncertainty in test genes relative to training genes was similarly observed in the GTEx dataset (Figure A12).

**Fig. 5:**
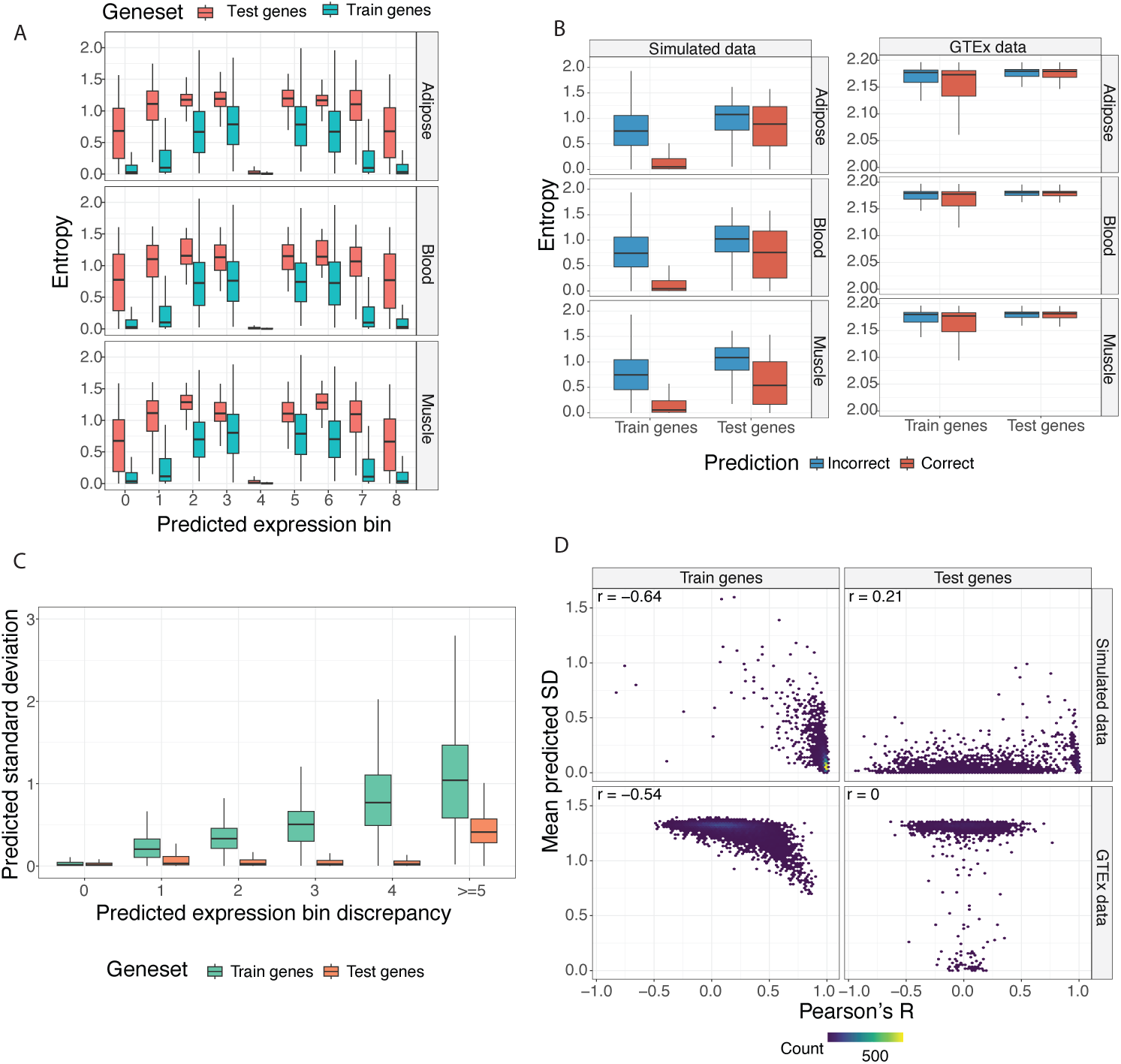
Predicted uncertainty reflects prediction errors in both simulated and GTEx data. (A) Predicted entropy varies across predicted bins in simulated data. Training and test genes follow similar trend but test genes have larger entropy. (B) Predicted entropy is larger for incorrect prediction (i.e. the predicted gene expression bin is different than the true gene expression bin) for simulated (left) and GTEx (right) data. Smaller difference in test genes compared to training genes. Red: correct predictions; Blue: incorrect predictions. Here we only show predictions exclude middle bin (i.e. we observed low entropy when the prediction is zero which can obscure the result). (C) Boxplots show the distribution of predicted standard deviation (Y-axis) versus the difference between predicted gene expression bin (the bin with thw largest predicted probability) and the true gene expression bin for training genes (green) and test genes (red) from simulated data. (D) Average predicted standard deviation per gene per tissue is negatively correlated with Pearson’s R for training genes but not in test genes. Y-axis is the mean predicted standard deviation per gene and X-axis is Pearson’s R, and their correlation is shown on the top left corner in each panel. Color represents the density of the points.

#### 2.4.3 Uncertainty as an indicator of prediction Correctness

Across both simulated and empirical data, predicted entropy was higher for incorrect predictions than for correct ones, though this difference was attenuated in test versus training genes and in GTEx versus simulations (Figure 5b). This pattern held true whether predictions in the central bin were excluded (Figure 5b) or included (Figure A12). Specifically, for central-bin predictions for training genes, correct classifications exhibited nearly zero uncertainty, whereas incorrect classifications generated high predicted entropy. However, this trend disappeared in the test gene set (Figure A12). Additional expression noise increased uncertainty. Training genes having mostly zero expression differences resulted in near-zero predicted values and correspondingly low entropy in simulations (Figure A13, A12) and in GTEx data, despite a lower baseline frequency of absolute zeros (10%) compared to the simulation (Figure A13). However, for test genes, the model frequently predicted a zero-difference state with falsely low entropy when the true expression difference was non-zero (Figure A13, A12). This illustrates a failure mode where the model defaults to a confident but incorrect zero prediction due to its inability to learn unseen eQTL effects.

In the simulation analysis, the predicted standard deviation for training genes strongly correlated with absolute prediction errors, validating the use of model-derived uncertainty as a proxy for prediction reliability (Figure 5c). Conversely, the model captured prediction error in test genes to a much lesser extent (Figure 5c). This discrepancy is further exemplified by comparing uncertainty against gene-level evaluation metrics. For both simulated and GTEx data, the mean predicted standard deviation per gene (aggregated across individuals and tissues) was negatively correlated with Pearson’s R for training genes, a relationship that dissolved for test genes (Figure 5d). Similarly, the mean predicted entropy demonstrated a significant negative correlation with prediction accuracy within the training gene set (Figure A14).

#### 2.4.4 Cross-run prediction stability as an alternate uncertainty proxy

Finally, we investigated whether variations in model predictions across independent training runs could serve as a more robust proxy for uncertainty in test genes. We compared predictions across distinct runs where models were trained on different donor cohorts but evaluated on an identical test donor set. For simulated data, only 2% of the predicted expression values diverged between runs. In contrast, 36% of the predictions varied in the GTEx data, revealing substantial donor-level heterogeneity (both genetic and non-genetic) in real tissue profiles that leads the model to capture distinct features depending on the training cohort. Notably, stable predictions (those identical across runs) achieved significantly higher accuracy than unstable ones in both settings (72% vs. 70% in simulation; 15% vs. 14% in real data; *χ*^2^ test *p <* 2*×*10*^−^*^16^ for both). Single-run predicted uncertainty closely mirrored this cross-run stability for test genes: both predicted entropy and standard deviation were minimized for predictions that were both stable and correct (Figure 6). However, the statistical divergence between the predicted probability distributions of different runs showed no meaningful correlation with gene-level performance metrics (Figure A15). Taking together, these results show that cross-run stability tracks the correctness of prediction to some extent, but cannot identify which unseen genes the model will predict correctly.

**Fig. 6:**
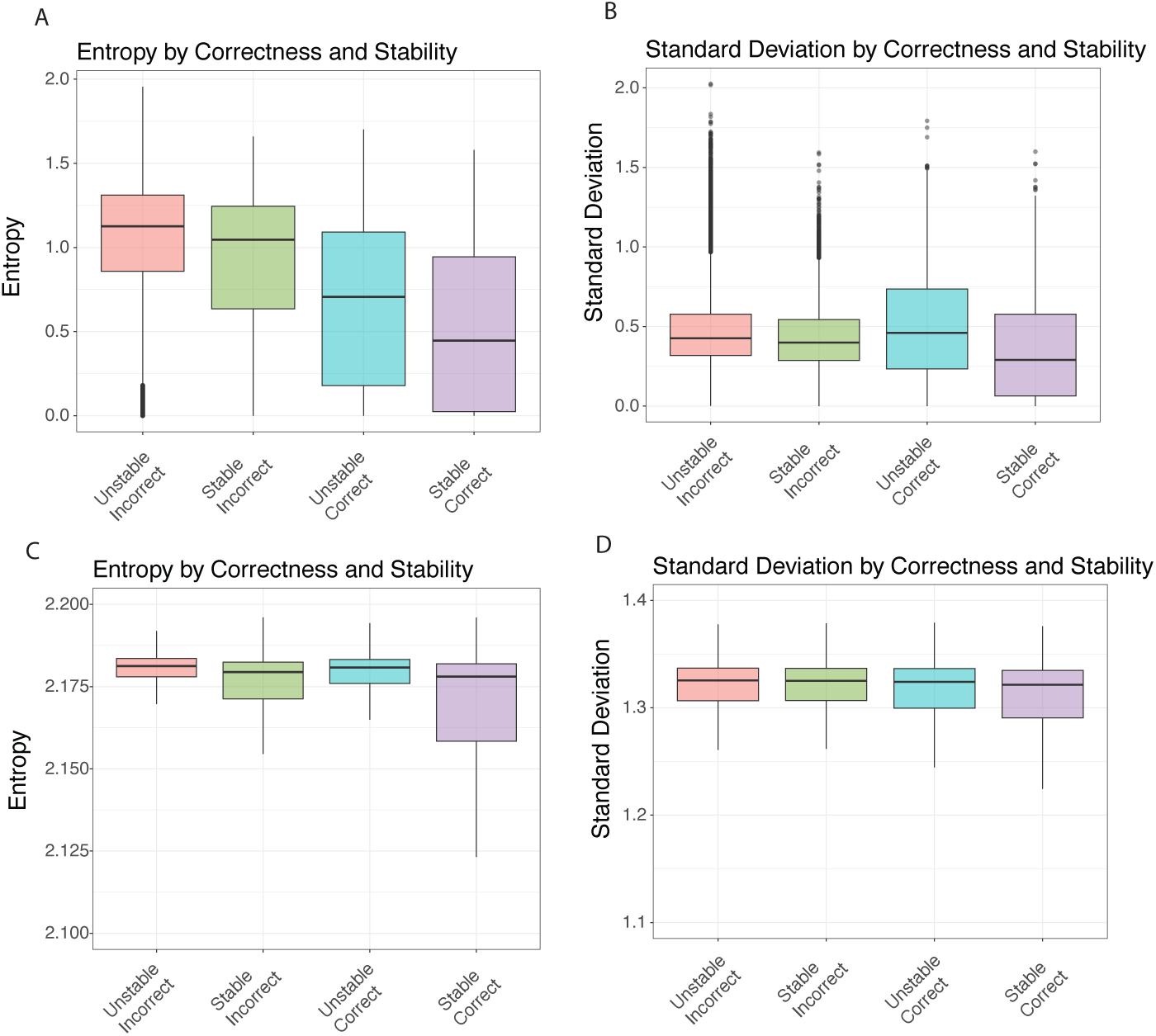
Predicted uncertainty reflects prediction correctness and prediction stability between the two runs in test genes in simulation (A-B) and real data (C-D). Predictions that are the same between the two runs (stable) have smaller predicted entropy and standard deviation than unstable ones. Stable and correct predictions have the smallest predicted entropy and standard deviation. Boxplots show the predicted entropy (A,C) and standard deviation (B,D) of the first run. Predictions near zero are excluded because of extreme small entropy and standard deviation in simulated data. Similar results for the second run, figure not shown.

## 3 Discussion

The quantitative benchmarks established in this study provide crucial insights into both the capabilities and the limitations of DNA sequence deep learning models for personalized genomics. Through the development and evaluation of CREAM, we have demonstrated that while modifications of neural network architecture in fine-tuning can successfully detect subtle cross-individual variations and uncover tissue-specific regulatory mechanisms, bridging the seen-to-unseen gene generalization gap remains a challenge.

Traditional S2F architectures face severe bottlenecks when adapting to individual genomes. Consecutive deep convolutional downsampling layers dilute fine-scale nucleotide polymorphisms within a nearly identical sequence background. CREAM effectively mitigates this structural limitation by creating direct links from intermediate convolutional layers to a customized cross-attention mechanism. This design draws architectural parallels to multi-scale hierarchical feature networks such as Feature Pyramid Networks (FPN) [21] and U-Net skip-connections [22], successfully isolating and preserving the multi-scale contexts of genomic bins with polymorphism. By combining this multi-scale feature retrieval with a contrastive learning framework focused explicitly on sequence differences between individual pairs, reminiscent of pairwise contrastive learning in computer vision [23] and genomics [24], CREAM significantly outperforms baseline architectures on training genes across both noise-free simulations and empirical GTEx transcriptomic data. Our architectural comparisons highlight that fully decoupling both the attention blocks and the downstream output layers across tissues (CREAM 2) is essential to accommodate divergent, tissue-specific regulatory profiles. This fully decoupled configuration successfully captured expected cross-tissue gene expression correlation.

Despite these structural enhancements, out-of-sample generalization to unseen test genes collapses to near-zero values across all evaluated methods, a result that mirrors broad generalizability limitations reported in recent deep sequence benchmarks [6]. This performance deficit indicates that current deep sequence models predominantly parameterize and memorize gene-specific regulatory contexts during their training phase rather than learning a universally transferable cis-regulatory mechanism. Our analysis of attention layers reveals a profound disconnect between variant identification and downstream quantitative expression prediction. For both training and test gene sets, CREAM’s architecture autonomously prioritizes fine-mapped causal eQTLs over background non-causal SNVs. When we introduced an auxiliary inductive bias during training, adding an auxiliary L1 loss function to minimize the distance between learned attention weights and known eQTL effect sizes, the model successfully detects causal eQTLs even within unseen test genes. However, this refined causal variant identification does not automatically map to superior gene expression prediction performance, which continues to fluctuate among genes. This points to highly complex genetic architectures and context-dependent regulatory mechanics that cannot be resolved simply by identifying causal eQTLs.

Another contribution of this work is the systematic characterization of predictive confidence through a hybrid discrete-continuous target formulation and both single- and cross-run metrics. While metrics within a single run like Shannon entropy and predicted variance serve as excellent proxies for prediction errors on training genes, this relationship breaks down for unseen genes. When evaluated on unseen test genes, the model regularly reverts to predicting a zero-difference expression state with falsely low entropy, even when the true expression difference is non-zero. Because the network cannot resolve the distinct gene regulatory architectures of unseen genes, it conservatively converges on a baseline prediction with high certainty.

To circumvent this limitation on unseen genes, we investigated cross-run prediction stability across independent initializations trained on varying donor cohorts. Stable predictions achieved significantly higher accuracy in both simulated and GTEx datasets, conceptually grounded in deep ensemble frameworks for uncertainty calibration [25]. However, the difference in accuracy between stable and unstable prediction is small, limiting the use of stability as a practical framework for prioritizing high-confidence variants.

To overcome training gene memorization and achieve generalization to unseen genes, one way is to utilize hierarchical structure to explicitly model distinct mechanistic stages like chromatin accessibility, splicing, and mRNA decay rather than mapping sequence to a single endpoint of gene expression. This multi-stages approach can potentially improve sequence-to-expression prediction, as current models successfully capture cross-individual variation in chromatin accessibility but not gene expression [26]. Models may also need to shift from cis-only paradigms to architectures that integrate trans-acting transcription and splicing factors directly. This cellular context is vital for generalizing localized regulatory grammar across unseen loci. Moreover, implementing self-supervised pre-training frameworks [27–29] allows networks to learn universal sequence syntax and evolutionary constraints. This pre-training pipeline could establish robust and transferable sequence representations that may be able to overcome training gene memorization.

## Code Availability

The CREAM source code and analysis scripts can be found in: https://github.com/xyz111131/CREAM/tree/main

## Acknowledgements

We thank Shiron Drusinsky and Sean Whalen for sharing the codes of Variformer, processed data and discussion. This work was supported by funding from the Biswas Family Foundation, the Keck Foundation, and the Dhaliwal Family.

## 5 Methods

### 5.1 Simulating gene expression

We obtained the identified eQTLs from SuSiE (https://github.com/eQTL-Catalogue/eQTL-Catalogue-resources/tree/master) based on GTEx V8 in blood, adipose and muscle (ftp://ftp.ebi.ac.uk/pub/databases/spot/eQTL/susie/QTS000015), as these tissues have more eQTLs and more individuals measured. We treated these eQTLs as ground truths and used the PIP * magnitude as coefficients to simulate gene expression using a linear model. Based on the individual genotype, we can simulate individual gene expression, then we standardize the gene expression across all individuals from GTEx V8.

### 5.2 Preprocessing GTEx dataset and model evaluation metrics

We utilized the GTEx V8 dataset for training and evaluation. We simulated individual genomes based on genotype data only keeping SNVs, similar to [6]. Our input genomic region is 49Kb around TSS of each gene. The length was chosen because of the memory contraint of our GPUs. The data partitioning and filtering workflow was structured as follows:

Donor partitioning: Individuals were split into training, validation, and test sets to evaluate cross-individual prediction performance through 10-fold cross validation. Due to computation cost, we only repeated the experiments on two partitions (fold 0 and fold 1). For each fold, we only kept donors with gene expression in all three tissues. For GTEx empirical data, we have 341 training, 47 valid, 42 test donors for fold 0 and 339 training, 46 valid, 45 test donors for fold 1. For simulated data, we have 670 training, 83 valid, 84 test donors for both folds. The real data has fewer donors because some of the donors do not have gene expression data.

Gene filtering and normalization: We restricted our analysis to variable genes by filtering for genes that met a minimum variance threshold (0.01 for simulated gene expression) and harbored at least one eQTL within 49Kb around TSS in at least one of the three analyzed tissues. We standardized gene expression for each tissue separately. For genes not expressed or measured in a particular tissue, we set the expression values to zero.

Gene partitioning: The remaining filtered genes were subsequently segregated into distinct training, validation, and test sets to robustly measure generalization to outof-sample genes. Training genes are on chromosomes 3, 6, 16, 19; validation genes are on chromosomes 6, 12, 18, 21; and test genes are on the remaining chromosomes. For simulated data, we have 4950 training, 1047 validation and 1646 test genes. For real data, we have 4529 training, 958 validation and 1538 test genes. The real data has fewer genes because some of the genes do not have gene expression data.

For evaluation, we converted predicted gene expression differences between individuals to absolute gene expression using linear regression, and computed Pearson’s r and *R*^2^ for each gene in training or test sets.

### 5.3 Model architecture and training

To model cross-individual gene expression variation from personalized genomes, we leverage the pre-trained Enformer architecture as a foundational sequence feature extractor, appending custom downstream attention and prediction heads designed to quantify variant-driven gene expression differences. The input to CREAM consists of a pair of genomic sequences from two individuals. Rather than evaluating absolute expression, the model predicts gene expression difference. The attention mechanism is designed to capture the interaction effects among SNVs and TSS, and the broader sequence background.

To achieve this, we extract multiscale features and construct a specialized cross-attention mechanism. Standard deep sequence models frequently suffer from a loss of fine-scale spatial resolution due to consecutive downsampling in convolutional layers. To preserve local variations, we systematically scan intermediate convolutional layers of Enformer to isolate the specific genomic bins that exhibit non-identical alleles between the two input genomes.

To construct the Query (*Q*) matrix, the embeddings from these polymorphic bins and those encompassing the TSS are concatenated across multiple convolutional layers and across both individuals. Concurrently, representations across all genomic bins are extracted from Enformer’s final attention layers and concatenated across both individuals to form the Key (*K*) and Value (*V*) matrices. We incorporated positional encodings into *Q* and *K* using Rotary Position Embedding (RoPE) [30]. This formulation allows the cross-attention layer to map the localized variant queries (*Q*) against the global regulatory environment (*K, V*), integrating distal elements that may modulate variant impact.

Following cross-attention transformation, an attention-pooling layer aggregates the spatially dispersed allelic signals across all polymorphic bins and the TSS into a unified, low-dimensional representation. A Multi-Layer Perceptron (MLP) then projects this pooled embedding to predict cross-individual gene expression differences. Crucially, the learned attention weights provide inherent model interpretability by quantifying the relative regulatory importance of each individual SNV. To assess biological fidelity, these empirical weights are directly benchmarked against Posterior Inclusion Probabilities (PIPs) derived from statistical fine-mapping via SuSiE.

Because eQTLs can manifest as either highly tissue-specific or broadly shared across human tissues, we developed three distinct architectural configurations of CREAM to evaluate multi-tissue expression dynamics:

CREAM 1 (Tissue-specific attention and shared output): To capture tissue-specific variant interactions while maintaining a unified prediction target, this model assigns distinct attention layers to each tissue but routes their output embeddings into shared downstream layers.

CREAM 2 (Fully decoupled attention and output): To allow complete parameter decoupling, this model fits each tissue with fully independent attention and downstream output layers.

CREAM 3 (Shared attention and tissue-specific output): To isolate tissue-agnostic regulatory features, this model uses a shared attention mechanism across all tissues, followed by independent tissue-specific output layers (2-layer MLPs) to predict distinct downstream expression levels.

For tissue-specific attention layer, we use 256 dimensions split into 2 heads, while we use 512 dimensions for the shared attention layer split into 4 heads.

During training and testing, the batch size is 20. We paired consecutive donors within a batch as input to CREAM model. During training, we also added random shifts of *±* 100 bp to the input sequences for data augmentation. Same shift is applied to all the individuals in a batch.

Training CREAM on 4 NVIDIA H100 80GB HBM3 GPUs for 20 epochs takes around 7 days for simulated data and 4 days for empirical GTEx data. We fine-tuned both parameters in the Enformer backbone and our custom downstream layers end-to-end. Optimization was performed using a learning rate of 1 *×* 10*^−^*^5^ by a OneCycleLR scheduler. To reduce sensitivity to expression outliers, we employed Smooth *L*_1_ loss between predicted and observed gene expression values. To incorporate known eQTLs as inductive biases into the attention mechanism, we introduced an auxiliary *L*_1_ loss function that minimizes the divergence between the learned attention weights and a prior target distribution derived from eQTL effect sizes. These effect sizes were normalized to sum to unity to match the scale of the attention weights. Prior to normalization, a pseudocount of 0.01 was added to the TSS coordinate to emphasize baseline attention on the core promoter while accommodating edge cases lacking active eQTLs. This regularizes the target vector by ensuring the model consistently prioritizes the core promoter region while safely handling edge cases where the sequence pair lacks active eQTLs. The final objective function is a combination of the primary prediction loss and this auxiliary guidance loss with equal weights.

### 5.4 Model ablation and baseline

To evaluate the benefit of extracting intermediate convolutional features from polymorphic bins, we constructed a “Contrast” model as an architectural ablation. Unlike CREAM, this model skips the fine-scale scanning of intermediate layers for non-identical sequence bins. Instead, its attention Query (*Q*) matrix is formed solely by concatenating the TSS-containing bin embeddings from both individuals. The Key (*K*) and Value (*V*) matrices are constructed using all spatial bin embeddings concatenating both individuals. The resulting cross-attention representations are then fed into a two-layer MLP with multi-task output heads to jointly predict cross-individual expression differences across multiple tissues.

As a standard benchmark, we implemented a Baseline model. This model extracts the Enformer sequence embedding exclusively from the bin encompassing the TSS. The TSS embedding is passed directly into a single linear projection layer to map the feature representations to individual gene expression across the target tissues without utilizing custom attention and contrastive mechanisms.

During training and testing, the batch size is still 20. Because of specific model architecture, we paired consecutive donors within a batch as input to Contrast model, while for Baseline model, we used each individual donor as input. The training schedule is similar as CREAM.

### 5.5 Uncertainty quantification and prediction stability

To assess whether CREAM can capture prediction confidence and distinguish true regulatory signals from background noise, we implemented an uncertainty quantification framework. This framework measures both single-model prediction confidence and multi-run ensemble stability. Rather than predicting raw continuous values alone, we reformulated the target as a hybrid discrete-continuous task to explicitly model a probability distribution over the outputs. The continuous, normalized cross-individual expression differences were discretized into 10 intervals (bins). To support this target structure, the network’s downstream projection head was modified to concurrently output two components: 1) categorical probabilities: a vector containing the probability of the expression difference falling into each of the 10 bins, generated via a softmax activation layer; 2) continuous local offsets: a continuous scalar value representing the predicted residual offset relative to the mean of each corresponding bin. Using the discrete-continuous hybrid output, we defined two statistical metrics to measure intrinsic prediction uncertainty: 1) Shannon entropy: computed over the categorical probability distribution to measure classification dispersion. A concentrated, unimodal distribution yields low entropy, indicating high model certainty, whereas a flat or multimodal distribution yields high entropy, reflecting a lack of model confidence; 2) prediction variance: measured across the continuous model predictions to track the expected numerical spread and variability of expression estimates.

To assess epistemic uncertainty, particularly when generalizing to unseen test genes, we evaluated prediction consistency across independent model initializations. We executed parallel training runs where models were optimized using different donor splits (cross-validation folds) but evaluated on an identical subset of test genes and test donors. We used two metrics to quantify cross-run alignment: 1) Jensen-Shannon Divergence (JSD): Computed between the predicted 10-bin categorical probability distributions of distinct runs to evaluate distributional consistency; 2) Prediction Stability: Categorized individual predictions as stable if independent runs predicted the exact same categorical bin, or unstable if the predicted bins diverged. These cross-run stability classifications and empirical divergences were subsequently benchmarked against true downstream prediction errors and performance metrics to determine if ensemble consistency serves as an accurate proxy for out-of-sample prediction accuracy.

## Declarations

The authors declare no competing interests.

## Appendix A Supplementary Figures

**Fig. A1:**
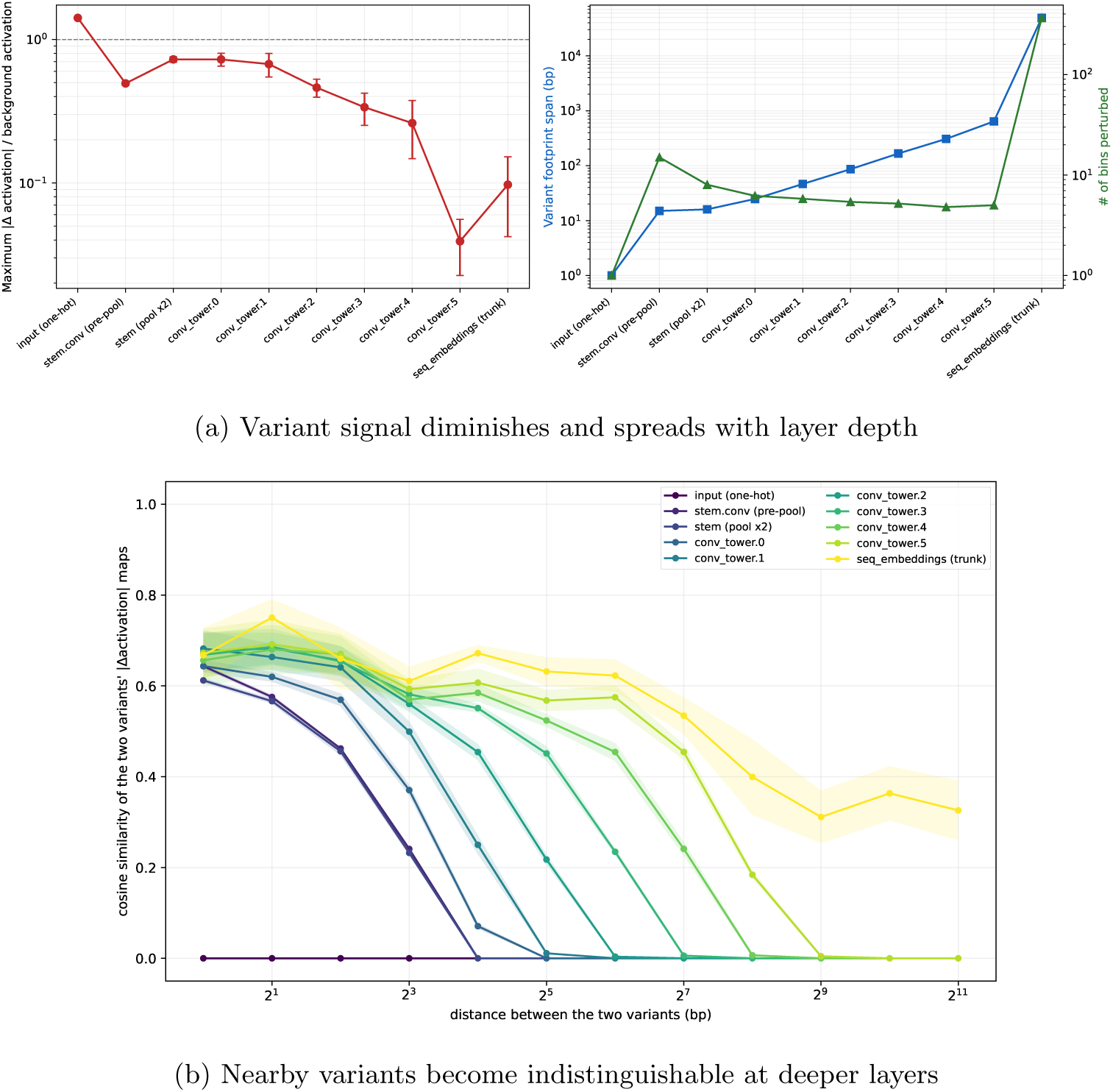
(a) Variant signal diminishes by Enformer’s downsampling convolution layers. Reference and single variant alternative sequences were passed through the frozen pretrained Enformer, and the per-bin absolute difference in activations was computed at consecutive layers of the downsampling convolution trunk and post-transformer embedding, whose resolution coarsens from 1 bp to 128 bp per bin. (Left) Maximum variant signal relative to background, defined as the maximum per-bin absolute difference in activations divided by the mean per-bin activation magnitude at that layer; error bars are *±* 1 SD across SNVs and the dashed line marks the background level. (Right) The SNV’s spatial footprint (span of bins carrying *>* 1% of the maximum difference, left axis, blue) and the number of bins perturbed (right axis, green) grow with depth, showing that the SNV signal is not only attenuated but also smeared across an increasingly wider region and almost constant number of bins at different convolution layers. These panels demonstrate that successive pooling both dilutes and delocalizes the effect of a SNV. (b) Nearby variants become indistinguishable at deeper layers, and the distance at which they merge grows with each downsampling step. For each reference context, an anchor SNV and a ladder of probe SNVs at increasing separations were introduced one at a time, and each variant’s effect map, Δ = *|act*(*alt*)*−act*(*ref*)*|*, was extracted at every layer and flatten i2n5side a genomic window (3072bp) centered at anchor. Each curve shows, for one layer, the cosine similarity between the anchor’s Δ map and the probe’s Δ map as a function of the genomic separation between the two variants (x-axis, log_2_ scale); curves are colored from shallow (dark) to deep (light) layers and show the mean over 100 randomly sampled TSS-centered intervals (hg38), with shaded bands giving *±* 1 SEM. At the input layer (darkest curve), the effect similarity of two distinct SNVs is almost 0 at every separation. As depth increases, nearby SNPs fall within the same pooled bin and receptive field, so their effect maps are similar for close-by variants. Thus the model loses the ability to resolve distinct polymorphisms that are closer together at deeper layers. All the curves show the mean over 100 randomly sampled TSS-centered intervals from Gencode v46 (hg38); the SNV was placed at a random position.

**Fig. A2:**
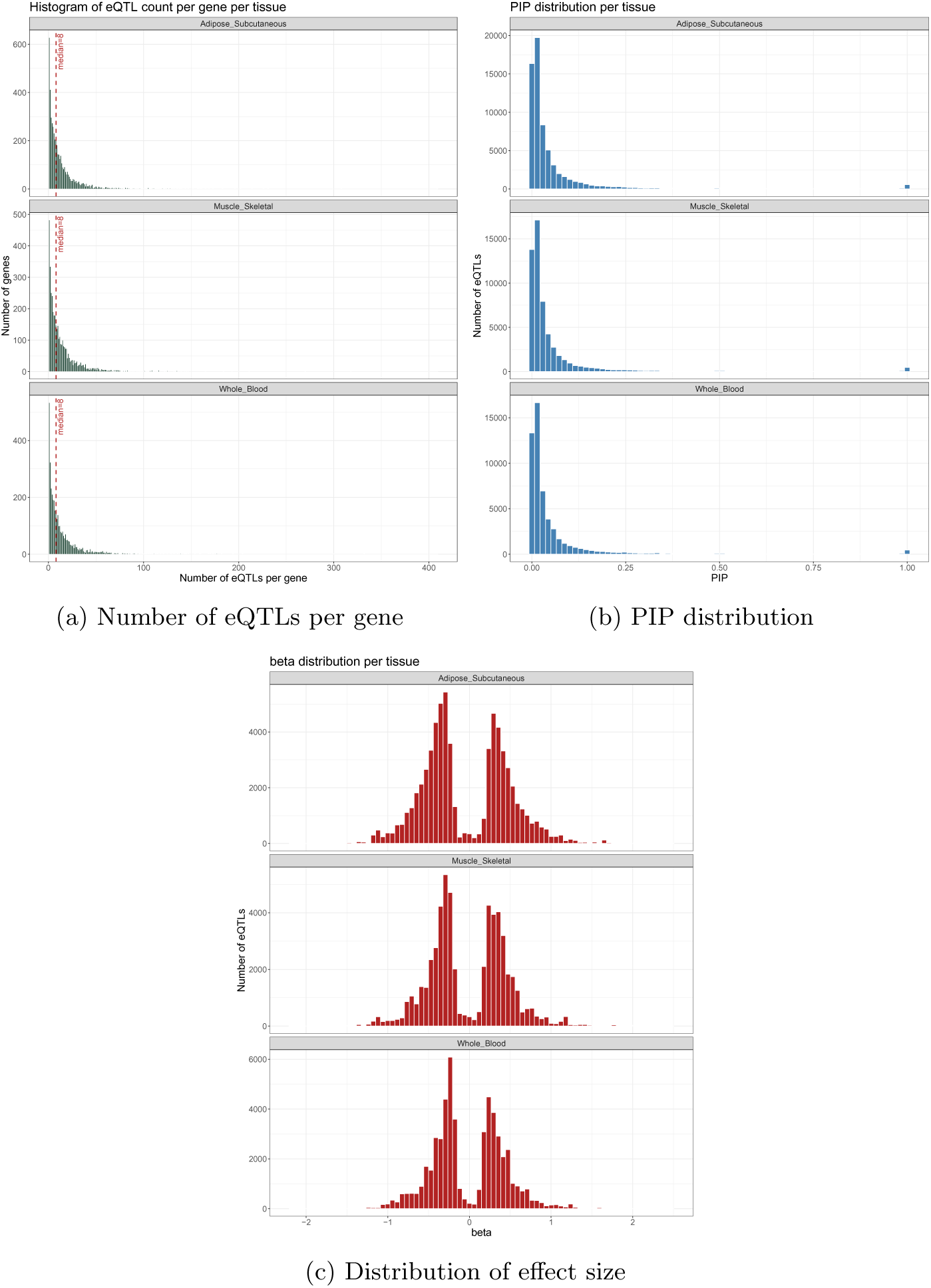
Summary statistics of eQTLs in blood, adipose and muscle. (a) Number of eQTLs per gene within 49kb around TSS. The median is 8 eQTLs per gene. (b) Distribution of PIP. Most eQTLs have PIP less than 0.2 while a few have PIP near 1. (c) Distribution of effect size of eQTLs. The effect size is far from zero but with in −1 and 1.

**Fig. A3:**
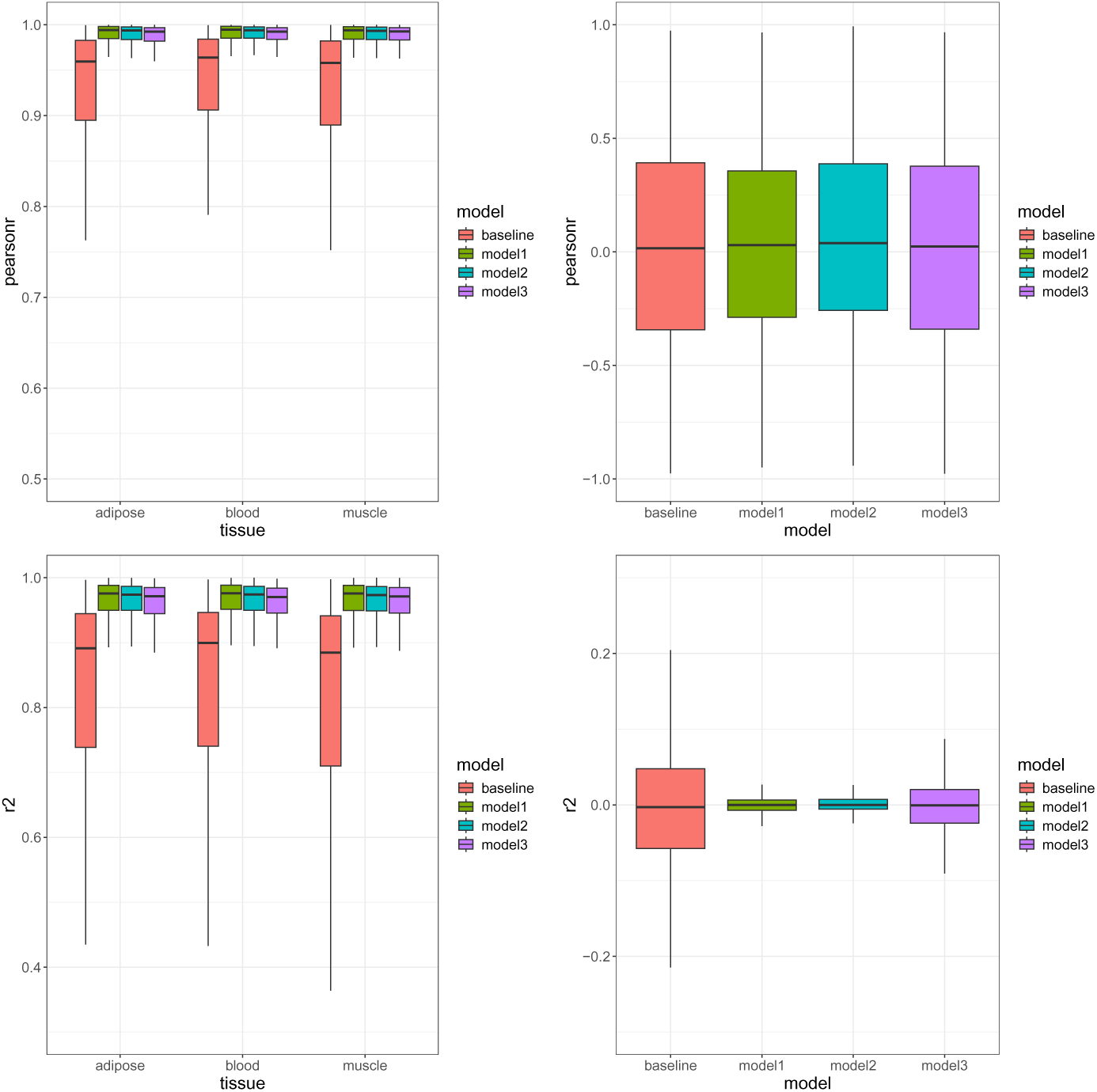
The boxplot shows the distribution of Pearson’s R and R2 (Y-axis) across training genes (left) and test genes (right) in different tissues (X-axis) using another train-test donor splits. Colors represent different models.

**Fig. A4:**
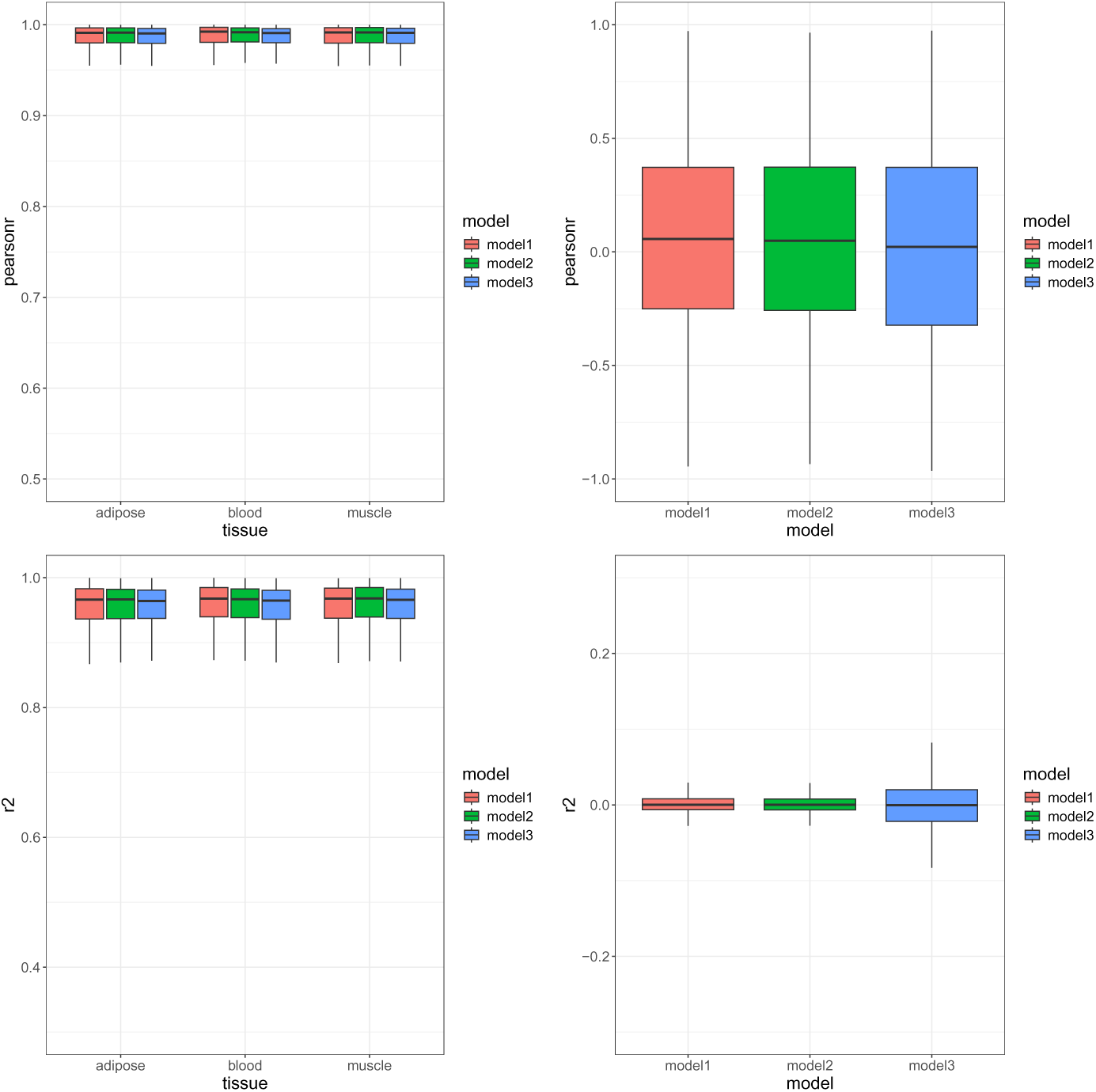
The boxplot shows the distribution of Pearson’s R and R2 (Y-axis) across training genes (left) and test genes (right) in different tissues (X-axis) using another random seed. Colors represent different models.

**Fig. A5:**
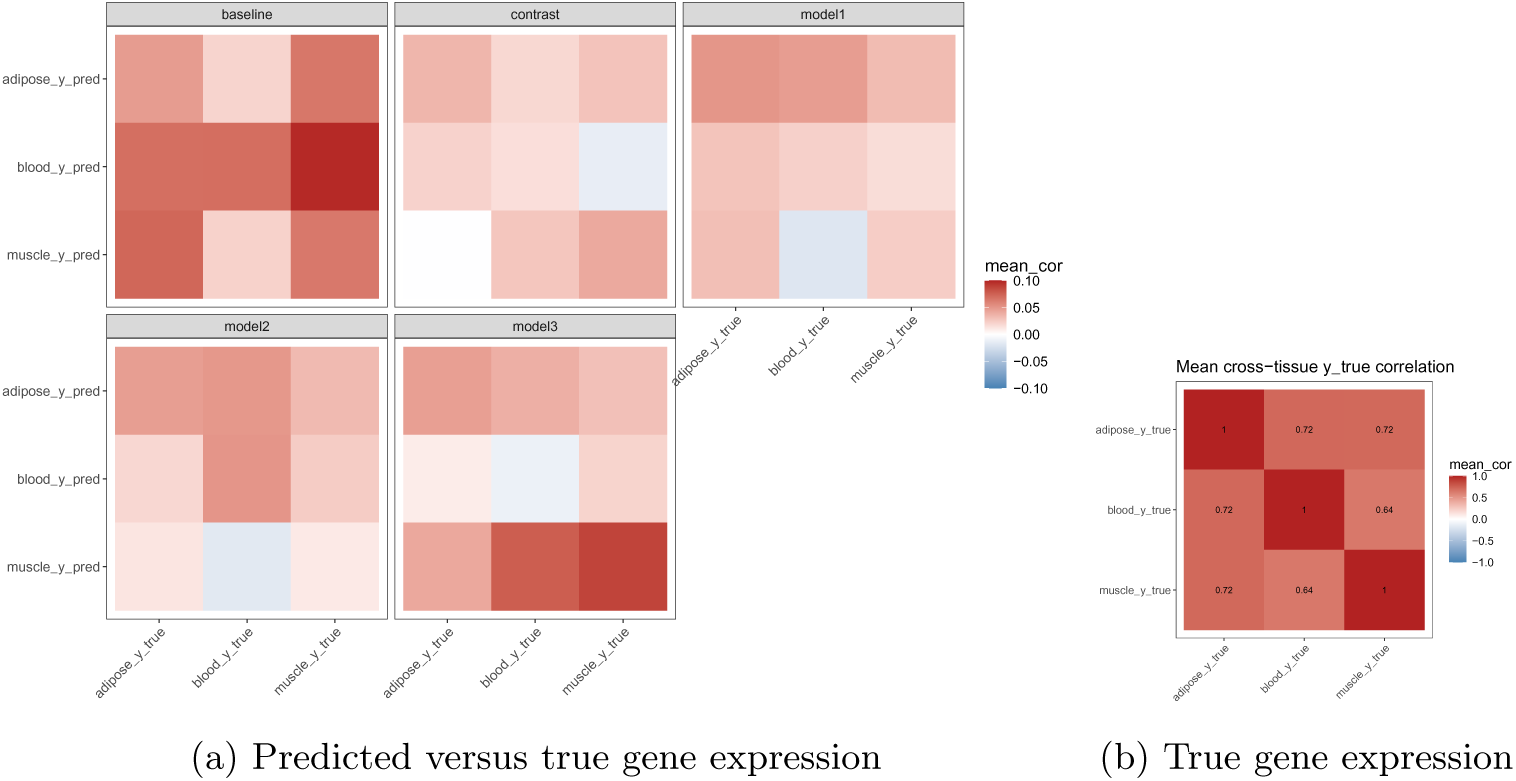
Cross-tissue correlation of simulated gene expression for tissue-specific genes. (a) Cross-tissue correlation of predicted and true gene expression in different models. Each row is predicted gene expression in different tissues and each column is true simulated gene expression. (b) Cross-tissue correlation of true simulated gene expression. Both rows and columns are true simulated gene expression. 366 tissue-specific genes within test gene set are selected as having at least one tissue-specific eQTLs with PIP *>* 0.5.

**Fig. A6:**
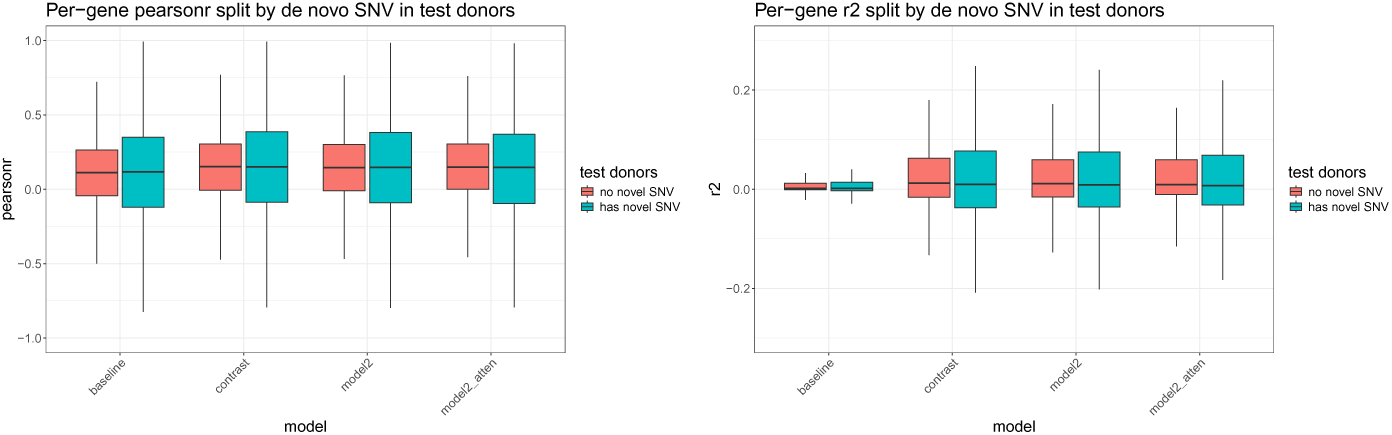
Cross-individual per-gene prediction performance is mostly unchanged for held-out donors carrying *de novo* SNVs. Per-gene prediction performance (Left: Pearson’s R, Right: R2) is shown across 4,529 training genes and three tissues (whole blood, skeletal muscle, subcutaneous adipose; pooled) for four models. Within each gene, donors were split by whether they carry a *de novo* SNV, i.e. a SNV in the gene’s 49kb TSS-centered input window absent from all training donors, and Pearson’s R or R2 were recomputed within each subgroup (subgroups less than 5 donors omitted).

**Fig. A7:**
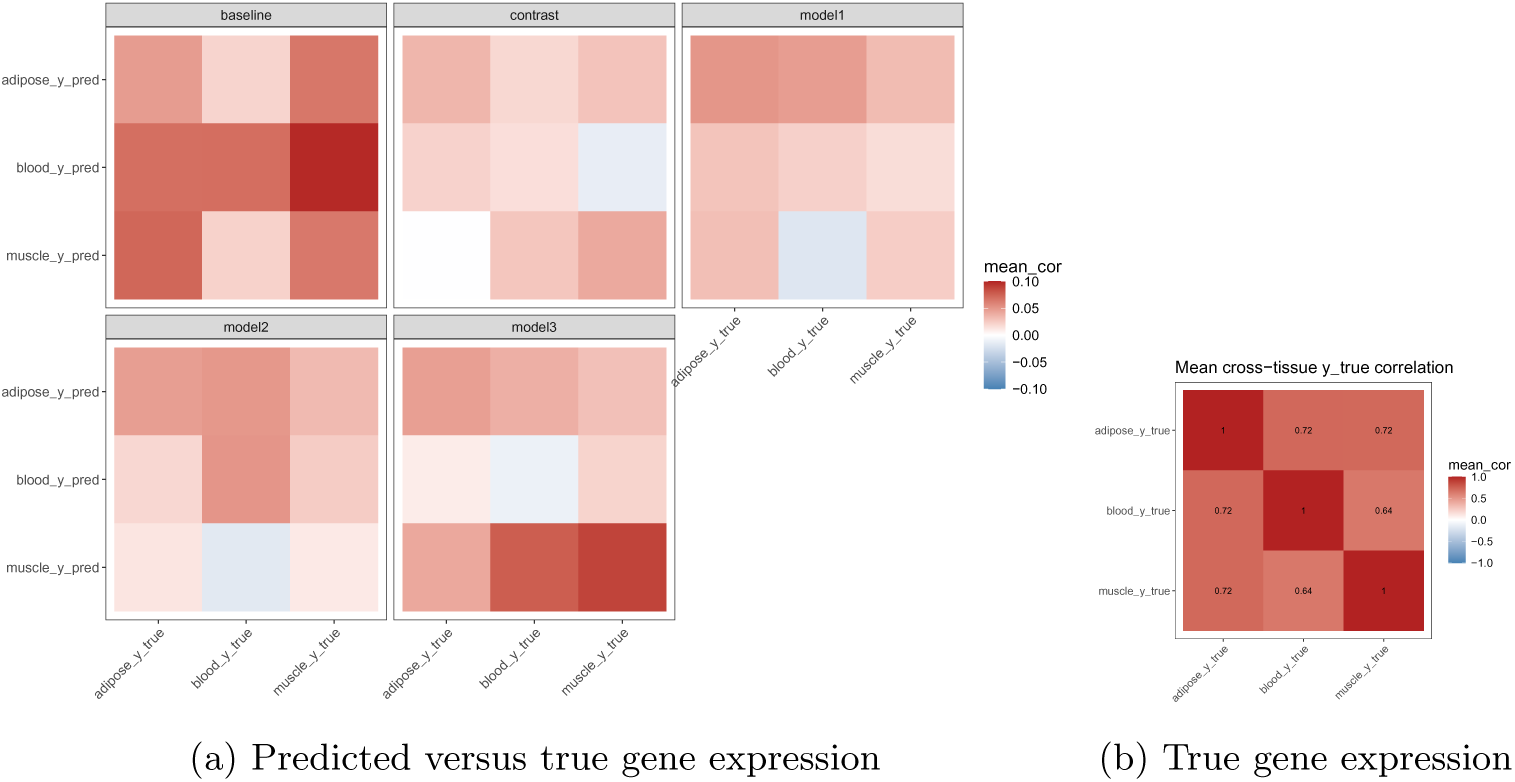
Cross-tissue correlation of gene expression for tissue-specific genes in GTEx data. (a) Cross-tissue correlation of predicted and true gene expression in different models. Each row is predicted gene expression in different tissues and each column is real gene expression. (b) Cross-tissue correlation of real gene expression. Both rows and columns are real gene expression. 1703 tissue-specific genes within training gene set are selected as having at least one tissue-specific eQTLs with PIP *>* 0.5.

**Fig. A8:**
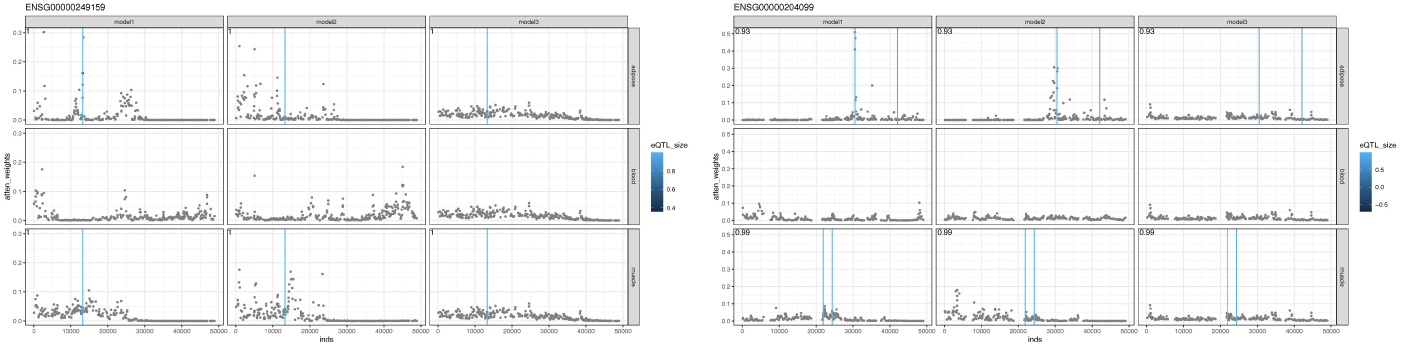
Manhattan plots show the average attention weights of each SNV in two example training genes in simulation. The attention weight is computed averaging over all test donor pairs and two runs with different seeds. Y-axis is the attention weights and X-axis is the input genomic coordinate. Each dot is a SNV colored by eQTL effect size. Vertical line indicates the eQTL with pip *>* 0.7. Pearson’s *R* is shown in the top-left corner. Calculation is restricted to the tissues with expression variation.

**Fig. A9:**
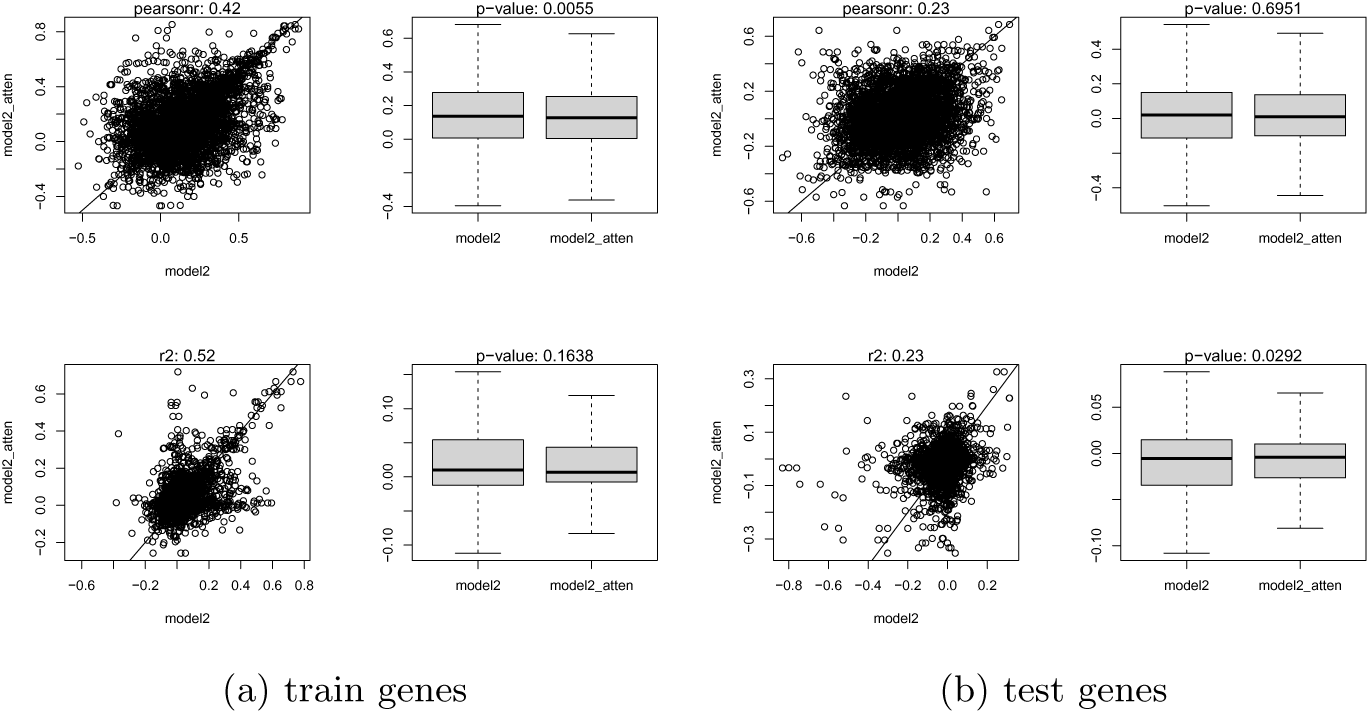
The scatterplot shows Pearson’s R and R2 between models with (X-axis) and without eQTL guided attention (Y-axis) for GTEx data. The boxplot shows the distribution of Pearson’s r and R2 (Y-axis) of these models (X-axix). Training genes (a) and test genes (b). Top: Pearson’s R; bottom: R2. The correlation of metrics between two models are shown on the top of each panel. p-values are obtained by paired two-sided Wilcoxon text.

**Fig. A10:**
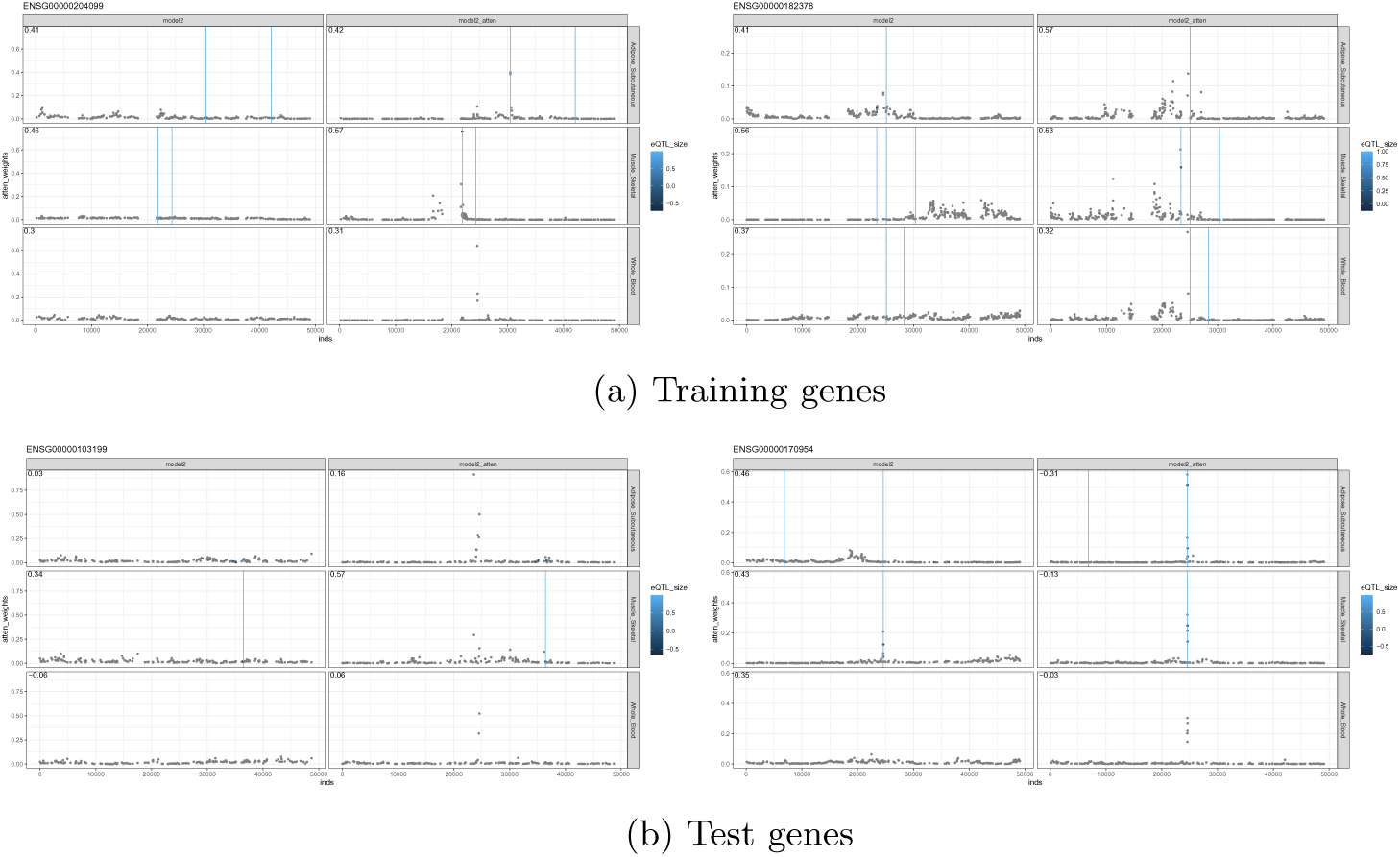
Manhattan plots show the average attention weights of each SNV in training (a) and test (b) genes in GTEx data. The attention weight is computed averaging over all test donor pairs. Pearson’s R of predicting gene expression is shown in the top left corner. Y-axis is the attention weights and X-axis is the input genomic coordinates. Each dot is a SNV colored by eQTL effect size. Vertical line indicates eQTL with PIP *>* 0.7. The two columns in each panel are original CREAM model (model 2) and eQTL guided attention model (model2_atten). eQTL guided attention model better captures causal eQTLs but the prediction performance varies. For ENSG00000170954 and ENSG00000182378, original CREAM model performs better or similarly than eQTL guided attention model; for ENSG00000103199 and ENSG00000204099, eQTL guided attention model performs better.

**Fig. A11:**
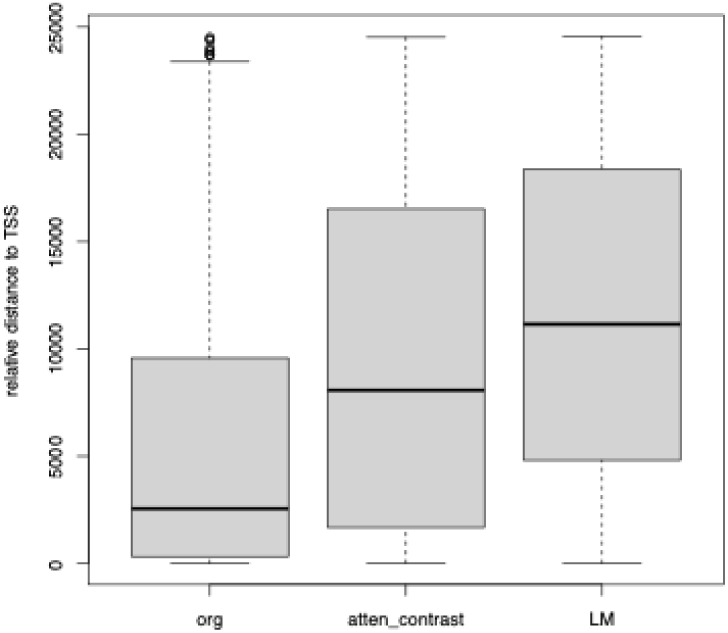
Distance to TSS of top 1000 high-score variants in 100 test genes by baseline, contrast and elastic net. For baseline and contrast model, high-score variants are computed by in silico mutagenesis, where for each SNV, the difference of gene expression with reference and alternative allele is computed. For elastic net, a penalized linear regression model is used to determine the coefficients of SNVs in predicting gene expression.

**Fig. A12:**
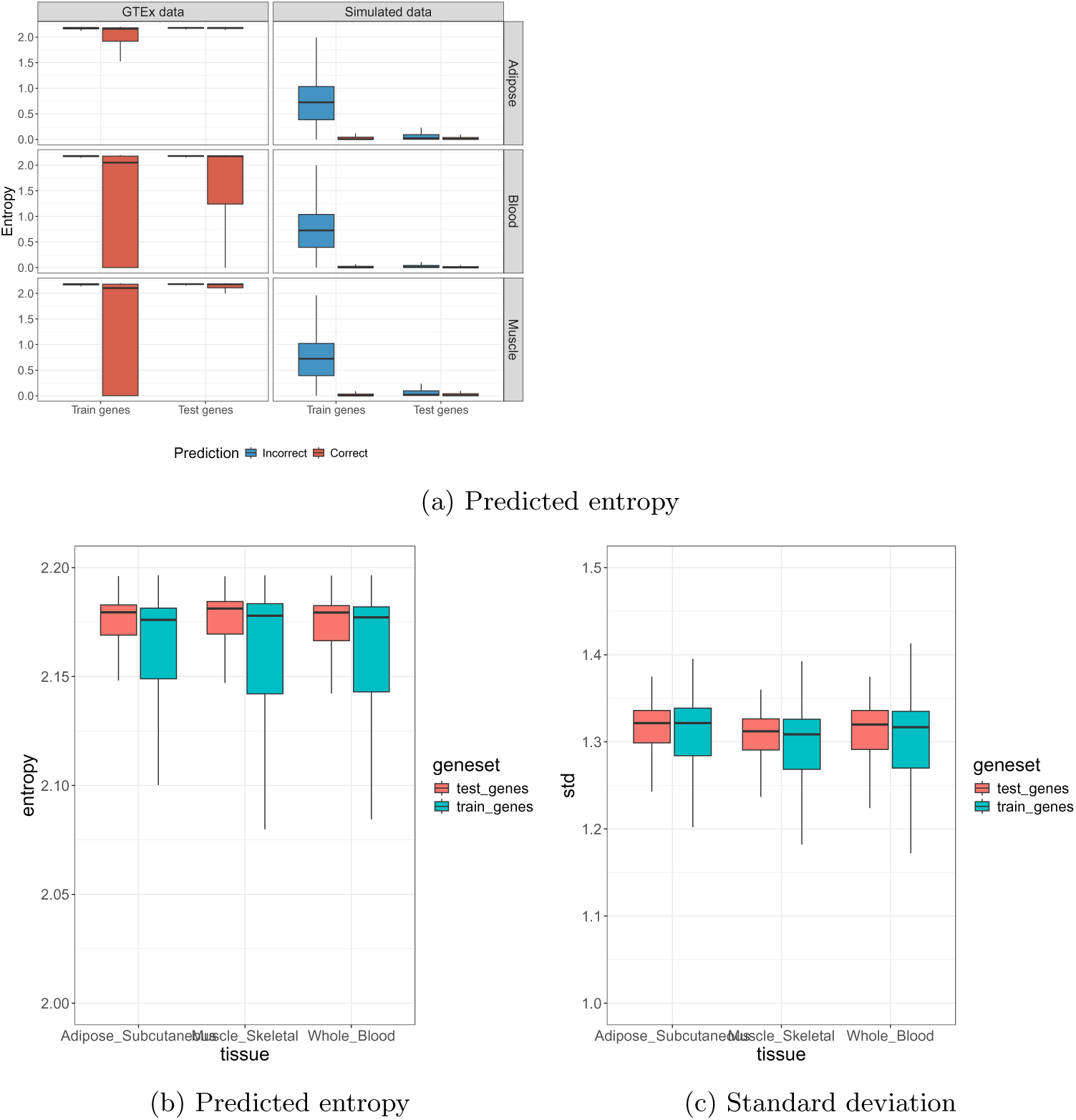
(a) Predicted entropy is larger for incorrect prediction (i.e. the predicted gene expression bin is different than the true gene expression bin) in GTEx and simulated data in multiple tissues. Red: incorrect predictions; blue: correct predictions. All predictions including middle bin. (b) Predicted entropy and (c) standard deviation are higher for test genes than train genes across tissues in GTEx data.

**Fig. A13:**
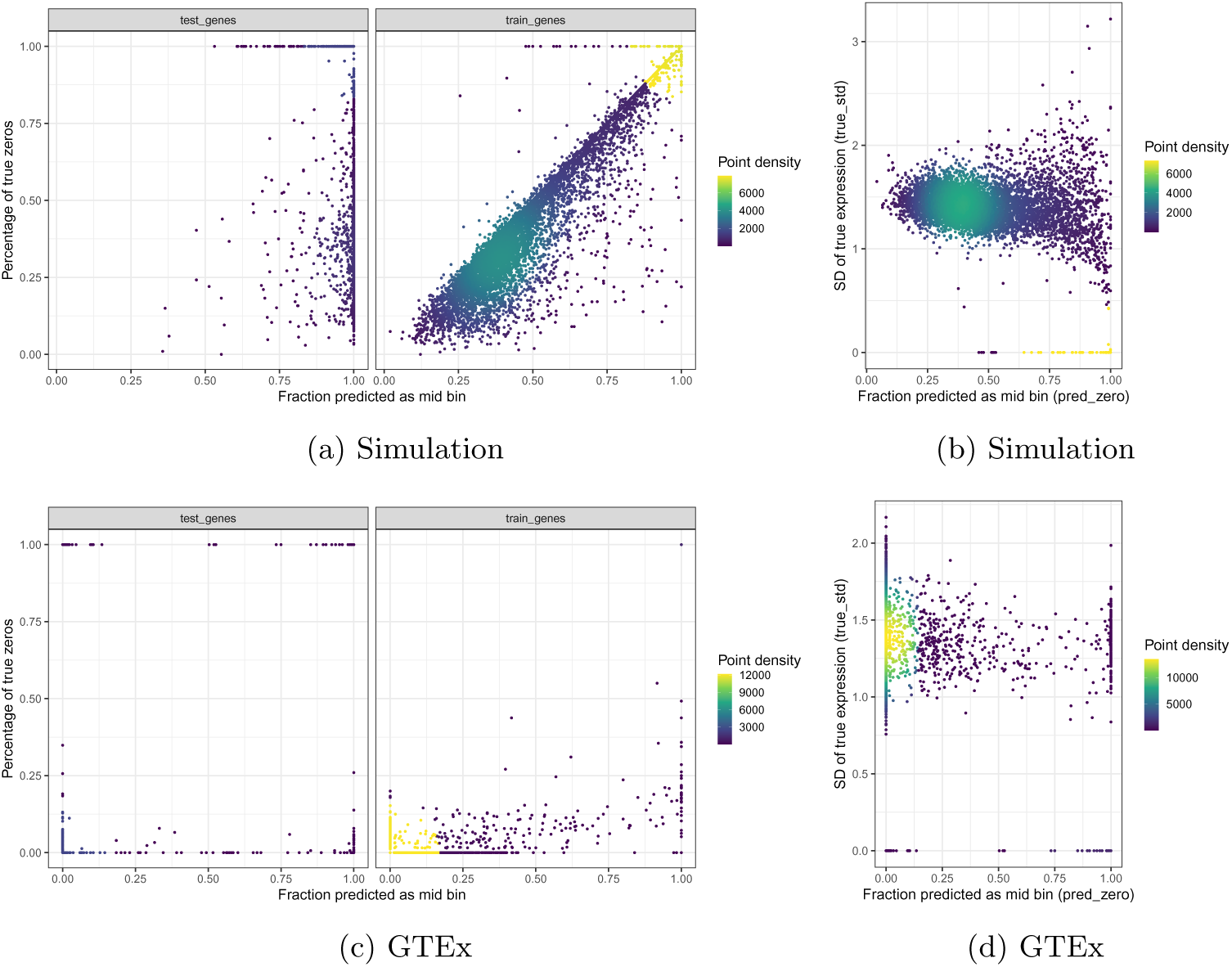
(a) and (c) Fraction of prediction in middle bin versus percentage of true zeros of gene expression difference in simulation and GTEx data. Fraction of prediction in middle bin is correlated with percentage of true zeros of gene expression difference in training but not in test genes. Percentage of true zeros of gene expression in simulated data is much larger than in real data. (b) and (d) Fraction of prediction in middle bin is anti-correlated with standard deviation of true gene expression difference in simulation and GTEx data. Each dot is a gene in a tissue. Although gene expression is normalized, some genes do not have eQTLs in certain tissue. Thus, their expression difference between donors are all zero and the predicted expression is also zero or close to zero. However, in test genes, many model predictions are zero but the true expression difference are not, as the model does not learn the eQTLs.

**Fig. A14:**
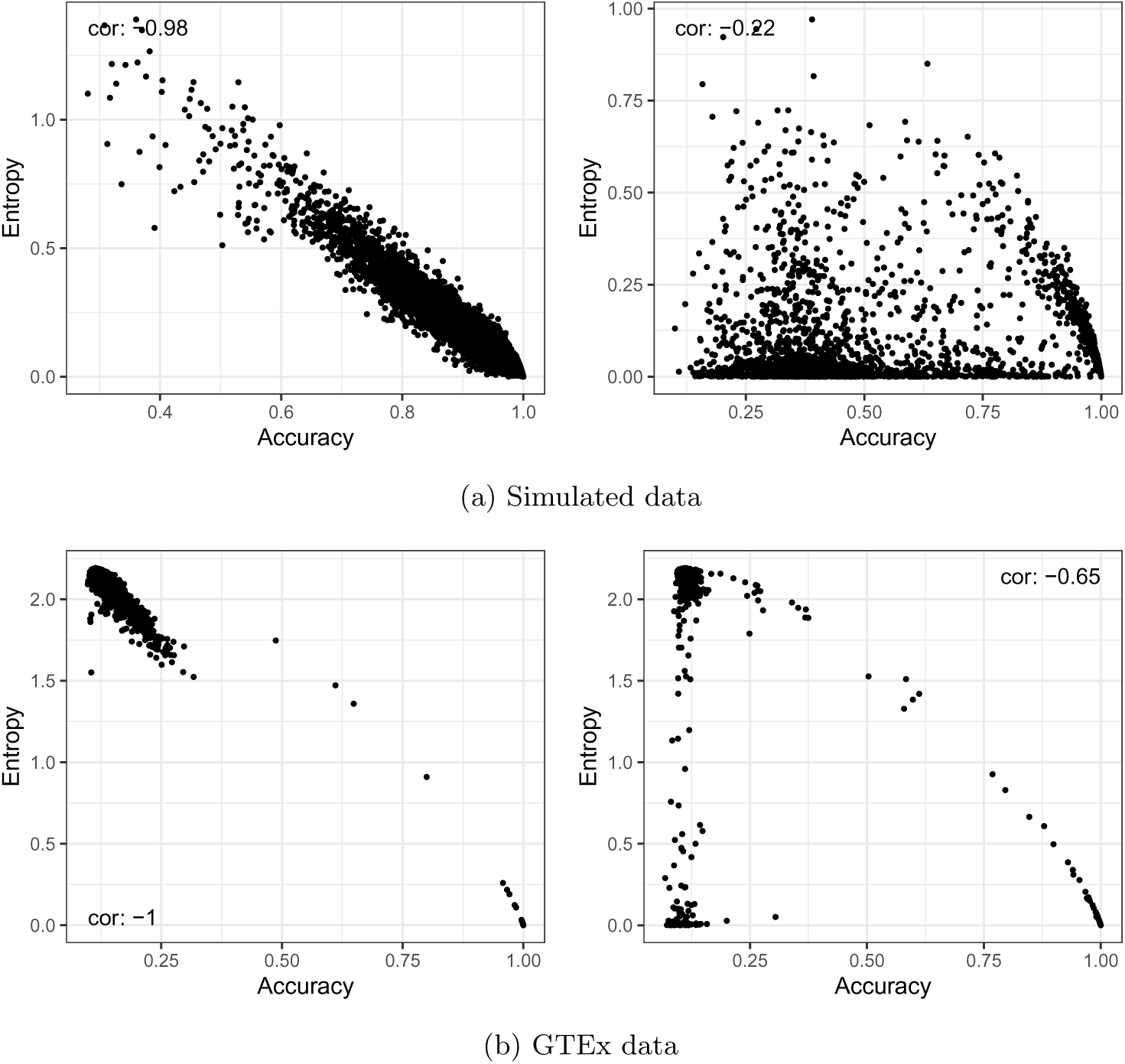
Average predicted entropy per gene per tissue is negatively correlated with prediction accuracy (i.e. whether the predicted expression bin is the truth). (a) Simulated data. (b) GTEx data. Left: training genes. Right: test genes.

**Fig. A15:**
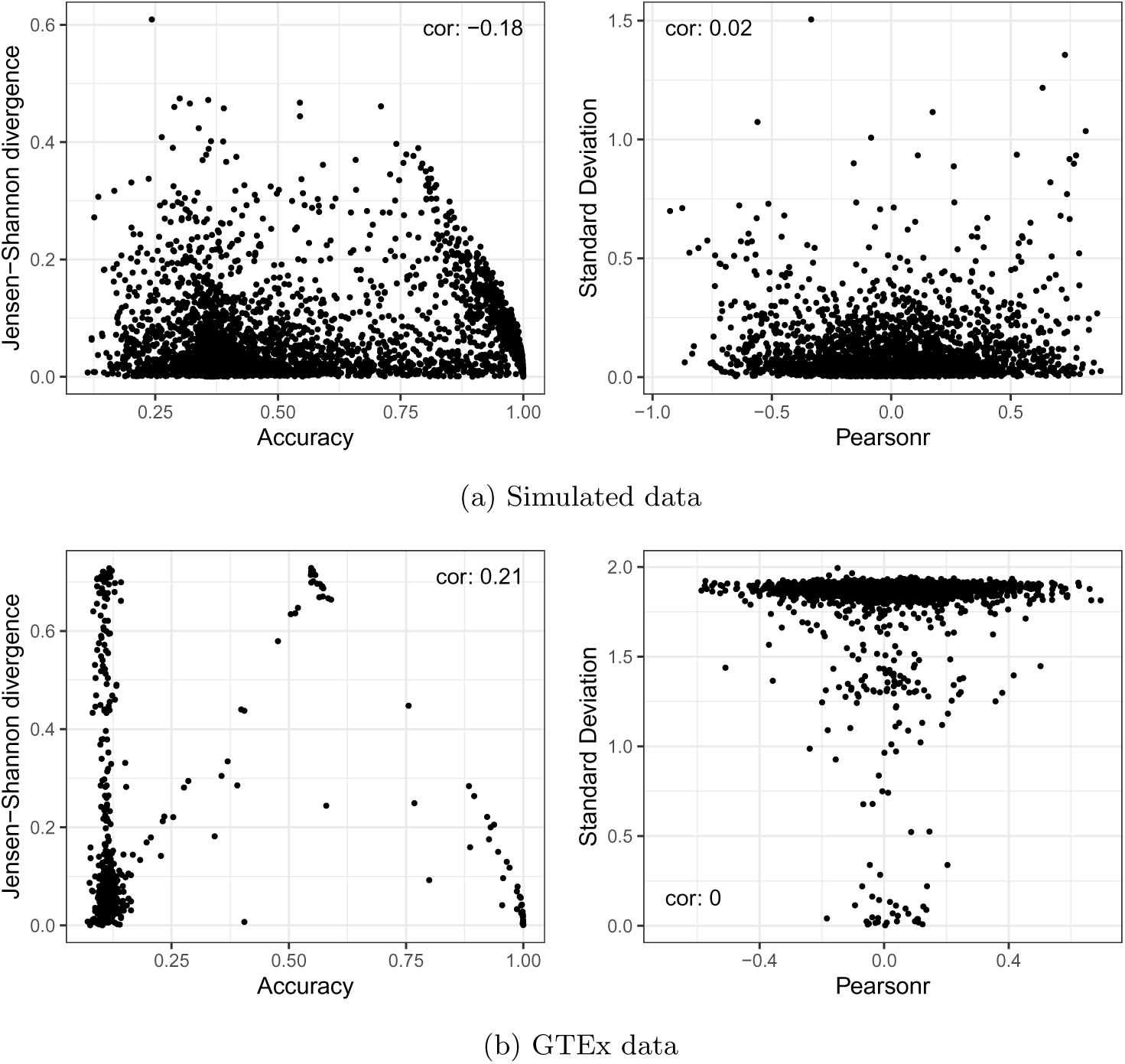
Prediction divergence between runs does not reflect prediction metrics in test genes in either (a) simulated data or (b) GTEx data. Average Jensen-Shannon divergence of predicted distribution between runs per gene per tissue is negatively correlated with prediction accuracy (i.e. whether the predicted expression bin is the truth) in simulation but not in real data. Average predicted overall standard deviation between runs per gene per tissue is not correlated with prediction accuracy.

